# Custom-molded headcases have limited efficacy in reducing head motion during naturalistic fMRI experiments

**DOI:** 10.1101/2020.03.27.012310

**Authors:** E Jolly, S Sadhukha, L.J. Chang

## Abstract

Effectively minimizing head motion continues to be a challenge for the collection of functional magnetic resonance imaging (fMRI) data. The use of individual-specific custom molded headcases is a promising solution to this issue, but there has been limited work to date. In the present work, we examine the efficacy of headcases in a larger group of participants engaged in naturalistic scanning paradigms including: long movie-watching scans (∼20-45min) and a recall task that involved talking aloud inside the MRI. Unlike previous work, we find that headcases do not reliably reduce motion during movie viewing compared to alternative methods such as foam pillows or foam pillows plus medical tape. Surprisingly, we also find that motion is *worse* when participants talk aloud while wearing headcases. These differences appear to be driven by large, brief rotations of the head as well as translations in the z-plane as participants speak. Smaller, constant head movements appear equivalent with or without headcases. The largest reductions in head motion are observable when participants were situated with both foam pillows and medical tape. Altogether, this work suggests that in a healthy adult population, custom-molded headcases may provide limited efficacy in reducing head motion beyond existing tools available to researchers. We hope this work can help improve the quality of custom headcases, motivate the investigation of additional solutions, and provide additional information about head motion in naturalistic contexts.

## 1. Introduction

Head motion is an ongoing problem in functional magnetic resonance neuroimaging (fMRI) research and has been estimated to account for 30-90% of the total signal variance (Friston et al., 1996). Notably, motion during scans produces highly variable signal disruptions that dramatically change the readout of the global and voxel signals (Jonathan D. Power et al., 2012; Satterthwaite et al., 2013; Van Dijk et al., 2012). Head motion is often exacerbated in developmental and clinical populations (Satterthwaite et al., 2012; Vanderwal et al., 2015) and makes it difficult to estimate task-specific activations when motion is correlated with stimulus onsets (Bullmore et al., 1999). Functional connectivity analyses (fcMRI) are particularly susceptible to head motion, which introduces spurious but systematic correlations across brain regions (Jonathan D. Power et al., 2012; Van Dijk et al., 2012; Yan et al., 2013). Spurious correlations demonstrate regional variability and are often more pronounced in prefrontal areas including the default-mode and prefrontal networks (Van Dijk et al., 2012; Yan et al., 2013).

A large body of work continues to investigate various post-acquisition preprocessing and analytic strategies to mitigate the effects of head motion on the fMRI signal. A non-exhaustive list of such strategies include: ICA-based nuisance removal, PCA-based nuisance regression, voxel-specific realignment regression (including Volterra expansion), scrubbing, white-matter and cerebrospinal fluid space nuisance regression, global signal regression, and multi-echo denoising (Behzadi et al., 2007; Friston et al., 1996; Hallquist et al., 2013; Jo et al., 2013; Kundu et al., 2017; Mowinckel et al., 2012; Muschelli et al., 2014; J. D. Power et al., 2018; Jonathan D. Power et al., 2012; Satterthwaite et al., 2012; Siegel et al., 2014; Tyszka et al., 2014; Yan et al., 2013) (for a review see: (Jonathan D. Power et al., 2015). However, another class of solutions involves *minimizing* head motion *during* the acquisition of brain volumes. While some technical solutions like prospective acquisition correction (Thesen et al., 2000) have been used to correct motion in near real-time, more common solutions involve situating individuals within the scanner in a way that prevents movement of the head from occuring in the first place. Such techniques include the use of foam head-coil stabilizers and a bite-bar, thermoplastic masks over the nose and brow, general foam padding packed into the head-coil, visual feedback systems that allow participants to adjust their head position during a study, and tactile feedback using medical tape (Bettinardi et al., 1991; Fitzsimmons et al., 1997; Krause et al., 2019; Menon et al., 1997; Jonathan D. Power et al., 2019; Thulborn, 1999; Zaitsev et al., 2015). Other techniques include training, mock-scanning, and using specialized movie stimuli to improve compliance in developmental and clinical populations (Vanderwal et al., 2019). While these approaches have proved promising, some make data collection more arduous and none completely solve the issue of data contamination by head motion. This has often led to poor widespread adoption of these techniques except in the case of developmental or clinical populations (Zaitsev et al., 2015).

A recent novel solution involves the use of custom-molded head stabilizers (“headcases”), developed on a per-individual basis and conformant to an individual’s unique anatomy. These custom head molds are milled from rigid styrofoam and distributed by the commercial company Caseforge (https://caseforge.co). Molds are produced from 3D optical scans of each participant’s head, providing a custom fit that accounts for the shape of each individual’s skull, neck, and facial structure. Taking such factors into account has been claimed to prevent motion and more precisely position participants within a scanner, while also increasing comfort. While advertised as a promising alternative to previous approaches for reducing motion, to date only a single systematic investigation into these claims has been published in the literature (Jonathan D. Power et al., 2019). One additional single participant dataset with and without headcases is also publicly available, but has been primarily used to investigate respiratory oscillations during multiband acquisition (Etzel & Braver, 2018). In their investigation, Power et al (2019) found that the use of headcases reduced motion during brief (4.8 min) resting state fMRI (rsfMRI) acquisitions in a group of 13 participants aged 7-28 years old. Their primary findings describe how headcases: (1) decreased motion in both rotational and translational axes, (2) reduced the fraction of scans with large movements measured using mean and median framewise displacement (a composite measure of head motion calculated from the realignment of functional volumes (Jonathan D. Power et al., 2012, 2015)), and (3) reduced the size of the small, constant movements throughout the scan.

In the present work, we test the efficacy of these same custom-molded headcases in the context of more “naturalistic” experimental designs that involve movie watching and talking aloud in the scanner environment. Many recent studies have utilized this “task-free” approach to probe neural representations and cognition because it can capture a larger degree of individual variability in cognition (Vanderwal et al., 2019). This approach also provides researchers with an opportunity to utilize rich datasets to ask a variety of questions, free of the constraints that come from pre-committing to a particular experimental design (Kriegeskorte et al., 2008). However, many of these studies often involve much longer acquisition times providing more opportunities for participants to move (Meissner et al., 2019), e.g. ∼15 minutes (Haxby et al., 2011) up to ∼45 minutes (L. J. Chang et al., 2018). Further, some studies ask participants to speak aloud during scanning, certainly exacerbating head motion, as participants must enunciate organically, while lying as still as possible (Baldassano et al., 2017; J. Chen et al., 2017; Silbert et al., 2014; Stephens et al., 2010; Zadbood et al., 2017). These designs provide an excellent opportunity to examine the efficacy of custom headcases in reducing motion under more demanding situations that increase the likelihood of movement.

Therefore, the present work builds upon the examination by Power et al (2019) in several key ways: a) we compare movement from datasets representative of recent naturalistic experiments using much longer acquisitions (∼45 minute continuous run); b) we report between group comparisons with larger sample sizes (*N* = 26-35 vs *N* = 13); c) we take advantage of between group comparisons that are matched on nearly every acquisition feature (i.e. scanner site, parameters, stimulus, etc.) or experimental task (i.e. active verbal recall of a previously watched movie); d) we compare movement from datasets in which participants are speaking aloud, providing a more rigorous test of the performance of headcases under more demanding scenarios. For consistency and direct comparison, we utilize the same approach as Power et al, focusing primarily on Framewise Displacement as a global metric for head motion (Jonathan D. Power et al., 2012, 2015) and temporal SNR (tSNR) as a metric for overall signal quality. In addition, we also examine differences and intersubject correlations in individual translation and rotational motion axes to test how headcases affect consistent shared motion by participants viewing the same movie.

Our principal analyses focus on between groups comparison while participants *viewed* a single episode of a television show or *talked* aloud about the narrative of that television show. Viewing comparisons comprise two datasets collected in an identical fashion at the same site, from the same population, viewing the same episode either with or without headcases. These comparisons also include an additional dataset collected in a similar fashion albeit at a different site, from a different but comparable population, using different acquisition parameters without headcases. This particular dataset served as an additional control to ensure that any differences (or lack thereof) between the two similarly acquired datasets were not driven by site, stimulus, or population idiosyncrasies. Talking comparisons comprise two of the three viewing datasets which were collected at different sites, but involved the same experimental task: freely recalling the narrative of each respective television episode with minimal time constraints. All datasets are described in Table 1.

**Table 1.** Description of datasets

| *Dataset* | *Site* | *Stimulus* | *Task* | *N* | *Headcase* |
| --- | --- | --- | --- | --- | --- |
| Dataset 1 | Dartmouth | Friday Night Lights | Viewing | 35 | No |
| Dataset 2 | Dartmouth | Friday Night Lights | Viewing | 26 | Yes |
|  | Dartmouth | Friday Night Lights | Talking | 26 | Yes |
| Dataset 3 | Princeton | Sherlock | Viewing | 17 | No |
|  | Princeton | Sherlock | Talking | 17 | No |

## 2. Methods

All reported analyses are comprised of observations from three different datasets. Datasets 1 and 3 come from previously published studies (L. J. Chang et al., 2018) and (J. Chen et al., 2017). A more detailed description of the data collection and preprocessing procedures are available in those initial publications, but we provide an abbreviated summary of the methods here. Separate manuscripts using Dataset 2 are forthcoming, but the subset of data used in the current manuscript was collected in a fashion identical to Dataset 1 with the addition of custom-molded headcases manufactured by Caseforge. Datasets 1 and 2 were collected at Dartmouth College while Dataset 3 was collected at Princeton University. All participants provided informed written consent in accordance with the experimental guidelines set by their respective institutions: the Committee for the Protection of Human Subjects at Dartmouth College and the Institutional Review Board at Princeton University.

### 2.1 Dataset 1 (FNL no-headcase)

#### 2.1.1 Participants & Procedure

Thirty-five (*M*_*age*_ *=* 19.0; *SD*_*age*_ = 1.07; 26 female) Dartmouth College undergraduate students were recruited from introductory psychology and neuroscience courses, participating for either monetary compensation ($20/hr) or for partial course credit. Each participant watched one episode of the television drama *Friday Night Lights* (FNL) while undergoing one continuous run of fMRI. Video was projected onto a rear-projection screen inside the magnet bore using an LCD projector and viewed with an angled mirror. The episode was displayed using the Psychopy toolbox in Python and synchronized to the onset of MRI data acquisition (Peirce et al., 2019). Audio was delivered using MR compatible in-ear headphones (Sensimetrics S14 https://www.sens.com/products/model-s14/). Participants were situated in the scanner with foam pillows and medical tape attached to their foreheads which provided tactile feedback (Krause et al., 2019) regarding head motion. Participants were also carefully instructed to lie as still as possible. For more details, see methods for Study 2 in (L. J. Chang et al., 2018).

#### 2.1.2 Imaging Acquisition

Data were acquired at the Dartmouth Brain Imaging Center (DBIC) on a 3T Siemens Magnetom Prisma scanner (Siemens, Erlangen, Germany) with a 32-channel phased-array head coil. Raw DICOM images were converted to NIfTI images and stored in the brain imaging data structure (BIDS) format using ReproIn from the ReproNim framework (K. J. Gorgolewski et al., 2016; Visconti di Oleggio Castello et al., 2018). Functional blood-oxygenation-level-dependent (BOLD) images were acquired in an interleaved fashion using gradient-echo echo-planar imaging with pre-scan normalization, fat suppression, an in-plane acceleration factor of two (i.e. GRAPPA 2), and no multiband (i.e. simultaneous multi-slice; SMS) acceleration: TR/TE: 2000/25ms, flip angle = 75°, resolution = 3mm^3^ isotropic voxels, matrix size = 80 x 80, FOV = 240 x 240mm^2^, 40 axial slices with full brain coverage and no gap, anterior-posterior phase encoding. Functional images were acquired in a single continuous run of 45.47 minutes (1364 TRs) which began and ended with 5 TRs of fixation.

### 2.2 Dataset 2 (FNL with headcase)

#### 2.2.1 Participants

Thirty-six (*M*_*age*_ *=* 22.77; *SD*_*age*_ *=* 4.73; 27 female) Dartmouth College undergraduate and graduate students were enrolled for a three-part study, participating for either monetary compensation ($20/hr) or for partial course credit. All reported data and analyses come from a subset of part-one and part-three of this study. One participant rescinded their desire to participate half-way through the first session and was consequently dropped from the dataset entirely. A total of 7 subjects were excluded from all reported analyses due to issues with their customized headcases: 4 participants’ reported extreme discomfort with their headcases during the first session, which resulted in no headcase use in subsequent sessions; 3 participants used only the front or back of their headcases along with additional foam padding due to discomfort. Two additional participants in this sample did not use headcases at all, but were situated in the scanner using foam pillows and medical tape as in Dataset 1. This resulted in a total of 26 (*M*_*age*_ = 22.92; *SD*_*age*_ = 4.96; 18 female) participants with headcases and two without headcases. These two participants were combined with participants from Dataset 1 for all *viewing* comparisons reported below, but were not included in any *talking* comparisons.

#### 2.2.2 Procedure

Across three experimental sessions that took place within approximately one week, participants watched the first four episodes of the television show *Friday Night Lights* and performed several memory tasks that involved talking aloud while undergoing fMRI. All reported analyses consist of motion estimates while *viewing* the first episode during session one, and *talking* about all four episodes during a spoken recall task in session three; session two (not reported, manuscript forthcoming) involved watching additional episodes of the same show. This recall task was similar in nature to the recall task used by Chen and colleagues (2017), in Dataset 3 (see below). Participants were asked to recall aloud the narrative events of all four episodes they had previously seen. They were given 1 minute to plan their responses and were asked to try to speak for a minimum of 10 minutes and a maximum of 30 minutes. This task was manually ended by experimenters when individuals verbally indicated they were finished, or automatically ended when the maximum recall time elapsed. Audio recordings were acquired using a MR-compatible microphone and recording system (Optoacoustic FOMRI III+ http://www.optoacoustics.com/medical/fomri-iii/features).

#### 2.2.3 Headcase production

Prior to coming in for the multi-part study, participants were asked to visit the lab to have their heads scanned with a handheld 3D scanner purchased from the CaseForge company. 360° scans of each participant’s head were acquired using procedures provided by CaseForge, which were identical to those reported by Power et al (2019). Participants with short hair wore a swim cap while being 3D photographed, as hair shape interfered with the quality and fit of the resulting case molds. For participants with longer hair, a fitted hood that extended down to the neck was used to reduce interference with photography. Images were uploaded to the Caseforge website which verified the quality of scans and subsequently shipped a two piece customized styrofoam mold consisting of a front and back half. Headcases were utilized in lieu of any additional padding within the head coil of the MRI machine during acquisition. As in Dataset 1, participants were instructed to lie as still as possible, particularly when speaking aloud. All headcases used in this dataset were manufactured in late 2017 through August 2018.

#### 2.2.4 Imaging Acquisition

Data were acquired at the Dartmouth Brain Imaging Center (DBIC) on a 3T Siemens Magnetom Prisma scanner (Siemens, Erlangen, Germany) with a 32-channel phased-array head coil using the same acquisition parameters as Dataset 1. Reported analyses come from a single continuous *viewing* run of 45.47 minutes (1364 TRs) which began and ended with 5 TRs of fixation and a variably ranged *talking* run (*M*_*Minutes*_ = 20.09; *SD*_*Minutes*_ = 6.52; *Min*_Minutes_ = 12.1; *Max*_*Minutes*_ = 31.2) which began with 5 TRs of fixation and ended with 15 TRs of fixation.

### 2.3 Dataset 3 (Sherlock)

#### 2.3.1 Participants & Procedure

Twenty-two participants (*M*_*age*_ = 20.8, 10 female) from the Princeton community were recruited for monetary compensation ($20/hr). Of this sample, 17 met inclusion criteria in the published sample by Chen et al (2017) and thus were used in all reported analyses. We direct readers to the aforementioned manuscript for full procedural details, but in brief: participants watched a 48-min segment of the BBC television series *Sherlock* and subsequently verbally recalled the narrative of the show aloud while undergoing fMRI. The episode was projected onto a rear-projection screen in the magnet bore using an LCD projector and viewed with an angled mirror. The episode was displayed using the Psychophysics Toolbox for MATLAB and synchronized to the onset of MRI data acquisition (Kleiner et al., 2007). During the recall task participants were instructed to talk for a minimum of 10 minutes, and were allowed to talk for as long as they wished. Experimenters manually ended the scanning run during the recall task based on verbal indication from participants. Participants were situated with foam padding and instructed to remain very still while viewing and speaking.

#### 2.3.2 Imaging Acquisition

Data were collected on a 3T full-body scanner (Siemens Skyra) with a 20-channel head coil. Functional images were acquired using a T2*-weighted echo-planar imaging (EPI) pulse sequence (TR 1500 ms, TE 28 ms, flip angle = 64°, whole-brain coverage 27 slices of 4 mm thickness, in-plane resolution 3 × 3 mm2, FOV 192 × 192 mm2), with ascending interleaved acquisition. Reported analyses come from two *viewing* runs of 23 and 25 minutes long and a variably ranged *talking* run (*M*_*Minutes*_ = 22.23; *SD*_*Minutes*_ = 8.62; *Min*_Minutes_ = 10.95; *Max*_*Minutes*_ = 44.15).

### 2.4 Motion Estimation and Comparison

#### 2.4.1 Framewise Displacement and motion parameters

For all three datasets, head position and motion was estimated using the FSL tool MCFLIRT (Jenkinson et al., 2002) in a two-pass procedure by first aligning each volume to the middle volume of the run, then computing a new mean volume, and then realigning each volume to this mean template. This yielded three translation and three rotation estimates at each volume. These parameters were used to calculate Framewise Displacement (FD) using the approach in Power et al. (2012) and implemented in nipype (K. Gorgolewski et al., 2011). This metric reflects the summation of the absolute-value backwards-differences of each parameter. Angular rotation parameters were converted to arc displacement using a 50mm radius prior to summation (Jonathan D. Power et al., 2012, 2015, 2019). For each participant, these motion time-series were used to calculate five summary statistics following the approach in Power et al (2019): mean and median displacement (FD_Mean_, FD_Median_), proportion of high-motion volumes that exceeded 0.3mm of displacement (Spike_Proportion_), and mean and median displacement excluding high-motion volumes (FD_MeanExcluded_, FD_MedianExcluded_). In Dataset 3, because participants viewed the stimulus across two separate runs, motion estimates were first calculated and summarized *within run* and the average of each pair of summary statistics was used for all analyses.

In order to account for different numbers of individuals in each dataset, all group comparisons were performed using permuted independent-groups non-equal variance t-tests implemented in Pymer4 (Jolly, 2018). First, for each comparison, a t-statistic was computed using Scipy (Jones et al., 2001--). Then, group labels (i.e. with or without headcase) were randomly shuffled while retaining the original group sizes and a new t-statistic was computed. This procedure was repeated 5,000 times to generate a null distribution of t-statistics. P-values were calculated by computing the number of permutations that were greater than or equal to the original t-statistic with adjustment to avoid non-zero p-values (Phipson & Smyth, 2010). Bootstrapped 95% confidence intervals around the mean difference between groups for each metric were also computed in Pymer4 by resampling with replacement from each group 5000 times while preserving the original group sizes. For all comparisons of individual motion parameters, p-values were corrected for multiple comparisons using a false discovery rate (FDR) of *q* = 0.05.

To further interrogate non-significant results, equivalence tests were performed using the two-one-sided-tests (TOST) procedure (Lakens et al., 2018; Schuirmann, 1981) implemented in the pingouin python statistics library (Vallat, 2018). This was performed to estimate whether any non-significant differences between groups fell within a predefined range of *practical equivalence*. In other words, non-significant group differences in motion alone do not provide information about practical differences between groups that may still be of interest to fMRI researchers, as small differences may still be detectable with large enough sample sizes. We chose the equivalence bounds of +/−0.05mm based upon the findings from Power et al (2019). In the *viewing* condition, comparisons were made between the FNL without headcases (Dataset 1) and FNL with headcases (Dataset 2) samples, while in the *talking* condition, comparisons were made between the Sherlock (Dataset 3) and FNL with headcases (Dataset 2) samples. For each comparison, one parametric, one-tailed, independent-groups, non-equal variance t-test were performed by shifting the mean of one sample by the equivalence bounds prior to running each comparison (Vallat, 2018). Consistent with statistical recommendations, the reported p-value reflects t-test against the upper equivalence bound of interest (headcases – non-headcases) (Lakens et al., 2018).

#### 2.4.2 Common motion across participants

To investigate whether participants exhibited common motion when *viewing* the same stimulus and how headcases affected this common motion, we computed the intersubject-correlation (ISC) (Hasson et al., 2004; Nastase et al., 2019) of each pair of participants’ Framewise Displacement time-series, separately in Datasets 1 and 2 (FNL without and with headcases respectively). We then performed a between groups comparison of the mean ISC using the mixed modeling approach recommended by Chen et al (G. Chen et al., 2017) implemented via Pymer4 (Jolly, 2018). Specifically, to account for the non-independence of pairwise ISC values, we doubled the observations and fit a model with separate random intercepts for each participant in a pair. Then we divide the estimated degrees of freedom in half to compute a p-value.

While Framewise Displacement is effective in capturing brief differences in motion, we also investigated whether linear or non-linear drifts were shared by participants and reduced by headcases. To do so we performed the same between groups ISC analysis separately for each non-differenced, raw motion parameter (six in total). However, this analysis assumes that participants move in the same way over time. It is also possible that participants move at the same time points during the movie, but in different ways. This idea forms the basis of functional alignment techniques like hyperalignment (Haxby et al., 2011) and the shared response model (SRM) (P.-H. (cameron) Chen et al., 2015). These techniques are typically used to align fMRI time-series across participants by learning a new *shared* feature space which abstracts away from voxel locations that may suffer from misalignment due to idiosyncrasies in anatomy not well accounted for by normalization. We used SRM to estimate transformation matrices that uniquely project each participant’s realignment parameters into six shared latent motion components and performed a between-groups ISC analysis separately for each component. This analysis examines whether headcases reduce motion in any direction not strictly reflected by the three translation and three rotation axes of the motion parameters, and possibly induced by the stimulus itself.

#### 2.4.3 Temporal Signal-to-Noise Ratio

While reducing head motion is the primary intended effect of equipment like headcases, the ultimate goal is to improve the overall signal quality of the collected data. To examine whether headcases impacted data quality, we performed a between groups comparison of temporal signal-to-noise ratio (tSNR) using Datasets 1 and 2 because they shared a common stimulus and acquisition parameters. We first preprocessed each participant’s time-series using the approach described in Chang et al (2018). Briefly, we used the realigned time-series to compute a mean image that was used to estimate linear coregistration parameters with each participant’s anatomical data. Then we used a non-linear normalization to project each participant’s anatomical data onto a 3mm MNI152 template. Finally we concatenated and performed these transformations in a single step. These analyses were performed using Advanced Normalization Tools (ANTs) (Avants et al., 2009) with a custom pipeline written in nipype (K. Gorgolewski et al., 2011). Finally, time-series were smoothed using a FWHM gaussian kernel of 6mm. tSNR was computed by dividing the mean of each voxel within a normalized gray matter mask by its standard deviation and a between groups parametric t-test was performed at each voxel using the nltools library (L. Chang et al., 2019). Initial thresholding was performed by controlling the voxel-wise false discovery rate (FDR) at *q* = 0.05. This produced no significant results so reported analyses visualize an uncorrected threshold of *p* < 0.001.

#### 2.4.4 Supplementary Analyses

We performed additional exploratory analyses examining the impact of headcases on: a) Framewise Displacement over time; b) the linear drift of Framewise Displacement; c) the probability of exhibiting a high motion TR with increasing scan length; d) detectable respiratory frequencies within the realignment parameter time-series. Overall we find that headcases had little to no impact on these metrics. These results and figures are available in the Supplementary Materials.

## 3. Results

### 3.1 Lack of overall motion reduction with headcases while viewing a movie

Based on previous findings (Jonathan D. Power et al., 2019), we expected to see reliable reductions in head motion when participants were fitted with custom headcases during fMRI scanning. However, while viewing the same stimulus (episode 1 of *Friday Night Lights)*, collected on the same scanner, at the same site, with the same acquisition parameters, we found no significant difference in FD_Mean_ (Fig 1 top row, left column, blue and pink bars). Participants with headcases moved equivalent amounts on average (*M* = 0.129; *SD* = 0.102) compared to participants situated with only foam pillows and medical tape (*M* = 0.113; *SD* = 0.045), *t* = −0.758, *p* = 0.494. This was also true when comparing FD_Median_, FNL with-case (*M* = 0.078; *SD* = 0.03), FNL no-case (*M* = 0.085; *SD* = 0.029), *t* = 0.857, *p* = .393, and the proportion of volumes with motion in excess of 0.3mm (Spike_Proportion_) FNL with-case (*M* = 0.06; *SD* = 0.075), FNL no-case (*M* = 0.045; *SD* = 0.048), *t* = −0.917, *p* = .363 (Figure 1 middle and bottom rows, left column, blue and pink bars).

**Figure 1.**
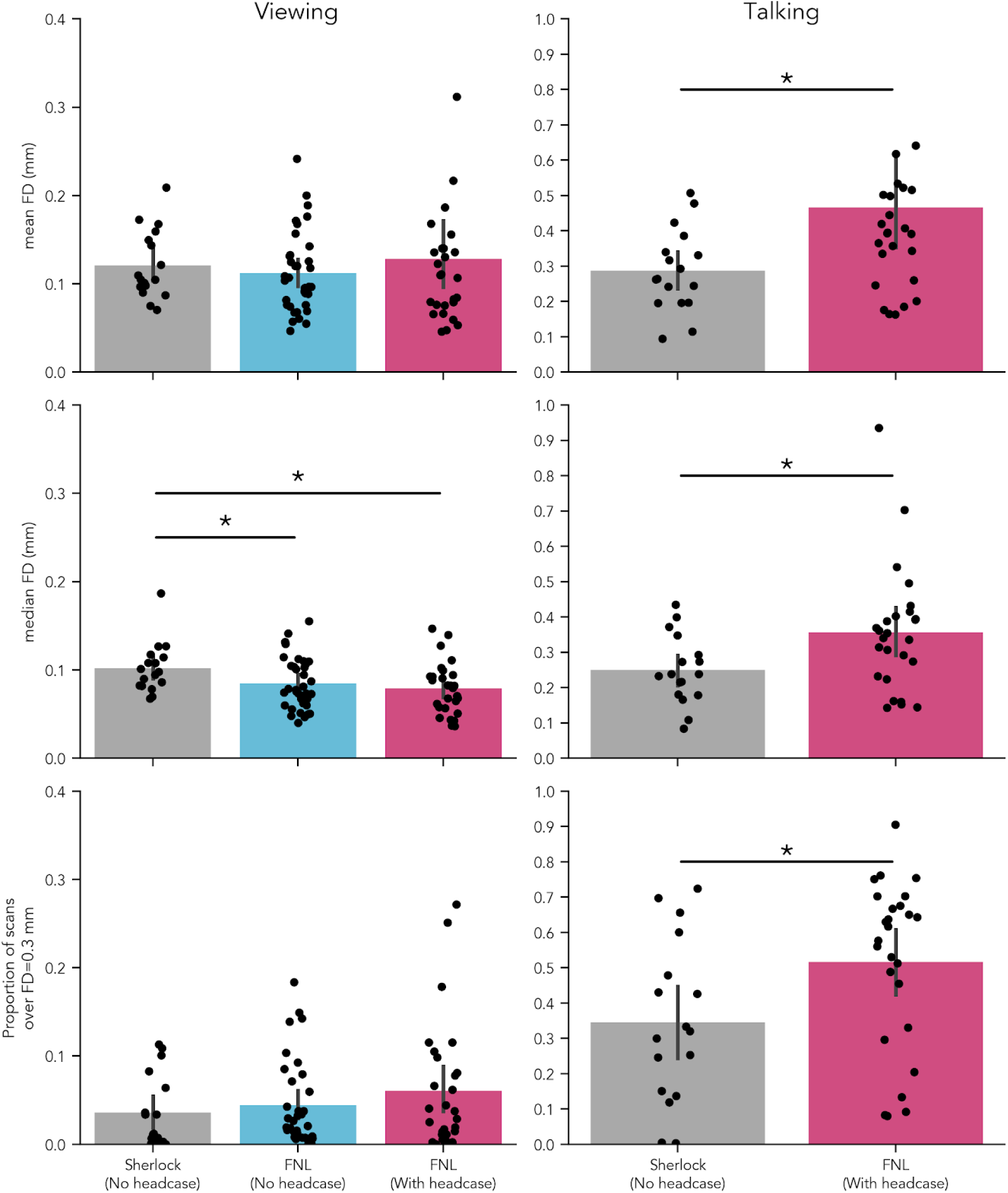
Headcase effects on Framewise Displacement when viewing and speaking inside the scanner. Top row: average of participant mean FD; middle row: average of participant median FD; bottom row: proportion of TRs in which FD exceeded 0.3mm. Blue and pink bars reflect two groups of participants who watched the same stimulus (Friday Night Lights) under the same data collection procedures (e.g. parameters, scanner site) except for the use of headcases. The grey bars reflect a group of subjects who watched a different stimulus (Sherlock) collected at a different site with different acquisition parameters. Sherlock participants, however, performed the same talking task inside the scanner albeit without headcases (right column). No significant improvement in mean or median FD or proportion of high motion TRs was observed with the use of headcases during viewing between participants watching the same stimulus. However, a significant difference was observed between the median FD of the Sherlock sample and both no headcase and headcase wearing FNL samples. During talking however, all metrics suggested a significant increase in motion while wearing headcases.

Despite being matched on nearly every dimension, we sought to ensure that our non-headcase sample exhibited motion typical of the range observed in similar naturalistic imaging studies, thus ensuring a fair comparison to our headcase sample. To do so, we compared motion estimates from our FNL non-headcase sample to another non-headcase dataset, in which individuals watched the first 48 minutes of the crime drama *Sherlock* (J. Chen et al., 2017) and were situated with foam pillows. Our non-headcase sample exhibited no significant differences in FD_Mean_ *t* = −0.703, *p* = .489, or Spike_Proportion_ *t* = 0.723, *p* = .466, but did exhibit a significantly lower FD_Median_, *t* = −2.061, *p* = .045. This translated to a significantly lower FD_Median_ for our *FNL headcase* sample relative to *Sherlock, t* = 2.60, *p* = .015. This suggests that motion in our non-headcase sample was comparable (and even slightly lower) relative to a similar existing non-headcase dataset. This also suggests that the lack of a significant difference between our non-headcase and headcase samples was unlikely due to something particularly unique about this dataset (Table 2).

**Table 2.** Head case comparisons while *viewing* movies

| *Comparison* | *Condition* | *Metric* | $M_{\text{Difference}}$ | $t$ | $p_{\text{perm}}$ |
| --- | --- | --- | --- | --- | --- |
| FNL no-case - FNL with-case | Viewing | FD<sub>Mean</sub> | -0.016<br>(-0.063 0.02) | -0.758 | 0.494 |
| FNL no-case - FNL with-case | Viewing | FD<sub>Median</sub> | 0.007<br>(-0.009 0.021) | 0.857 | 0.393 |
| FNL no-case - FNL with-case | Viewing | Spike<sub>Proportion</sub> | -0.015<br>(-0.049 0.015) | -0.917 | 0.363 |
| FNL no-case - Sherlock | Viewing | FD<sub>Mean</sub> | -0.008<br>(-0.032 0.015) | -0.703 | 0.489 |
| FNL no-case - Sherlock | Viewing | FD<sub>Median</sub> | -0.017<br>(-0.034 -0.002) | -2.061 | **0.045** |
| FNL no-case - Sherlock | Viewing | Spike<sub>Proportion</sub> | 0.009<br>(-0.016 0.033) | 0.723 | 0.466 |
| Sherlock - FNL with-case | Viewing | FD<sub>Mean</sub> | -0.008<br>(-0.053 0.031) | -0.349 | 0.776 |
| Sherlock - FNL with-case | Viewing | FD<sub>Median</sub> | 0.024<br>(0.007 0.042) | 2.6 | **0.015** |
| Sherlock - FNL with-case | Viewing | Spike<sub>Proportion</sub> | -0.024<br>(-0.06 0.009) | -1.375 | 0.185 |

However, because a lack of statistically significant differences does not necessarily provide evidence for the null hypothesis, we instead tried to better quantify these null results using equivalence testing (Schuirmann, 1981). Using the TOST procedure, we defined +/−0.05mm as the *equivalence bounds* of our comparisons, i.e. a range of mean-differences in head motion that fMRI researchers may consider “statistically equivalent” in (Lakens et al., 2018). This range was set based upon the reported *within-participant* average improvement in FD_Mean_ by Power et al (2019). We found that the observed mean differences and bootstrapped 95% confidence intervals for all motion metrics between headcase and non-headcase participants participants fell within this equivalence range: FD_Mean_ *p* = 0.002^1^; FD_Median_ *p* < .001 (Fig 5, left column, blue points). Together, these findings suggest that headcases provide limited efficacy in reducing overall head motion in longer scanning scenarios where participants have more opportunities to move.

### 3.2 Lack of small motion reduction with headcases while viewing a movie

Following the approach of Power et al (2019), we repeated the previous analysis, this time excluding high-motion volumes per individual (FD > 0.3mm) prior to computing and comparing summary statistics. This analysis assesses the efficacy of headcases in reducing smaller head movements by ignoring parts of each individual’s time-series that contain substantial motion. Our findings are largely similar to the previous results. We found no significant differences in FD_MeanExcluded_ (Fig 2 top row, left column, blue and pink bars) or FD_MedianExcluded_ when comparing participants with and without headcases viewing the same stimulus. The FD_Mean_ of participants with headcases was equivalent on average (*M* = 0.085; *SD* = 0.026) compared to participants situated with only foam pillows and headtape (*M* = 0.093; *SD* = 0.025), *t* = 1.186, *p* = .244. This was also true of FD_Median_, FNL with-case (*M* = 0.073; *SD* = 0.025), FNL no-case (*M* = 0.081; *SD* = 0.025), *t* = 1.33, *p* = .192. Equivalence tests using the same range as before also suggested that observed differences in FD_MeanExcluded_ and FD_MedianExcluded_ were of practical equivalence, *ps* < .001 (Fig 5, left panel, red points).

**Figure 2.**
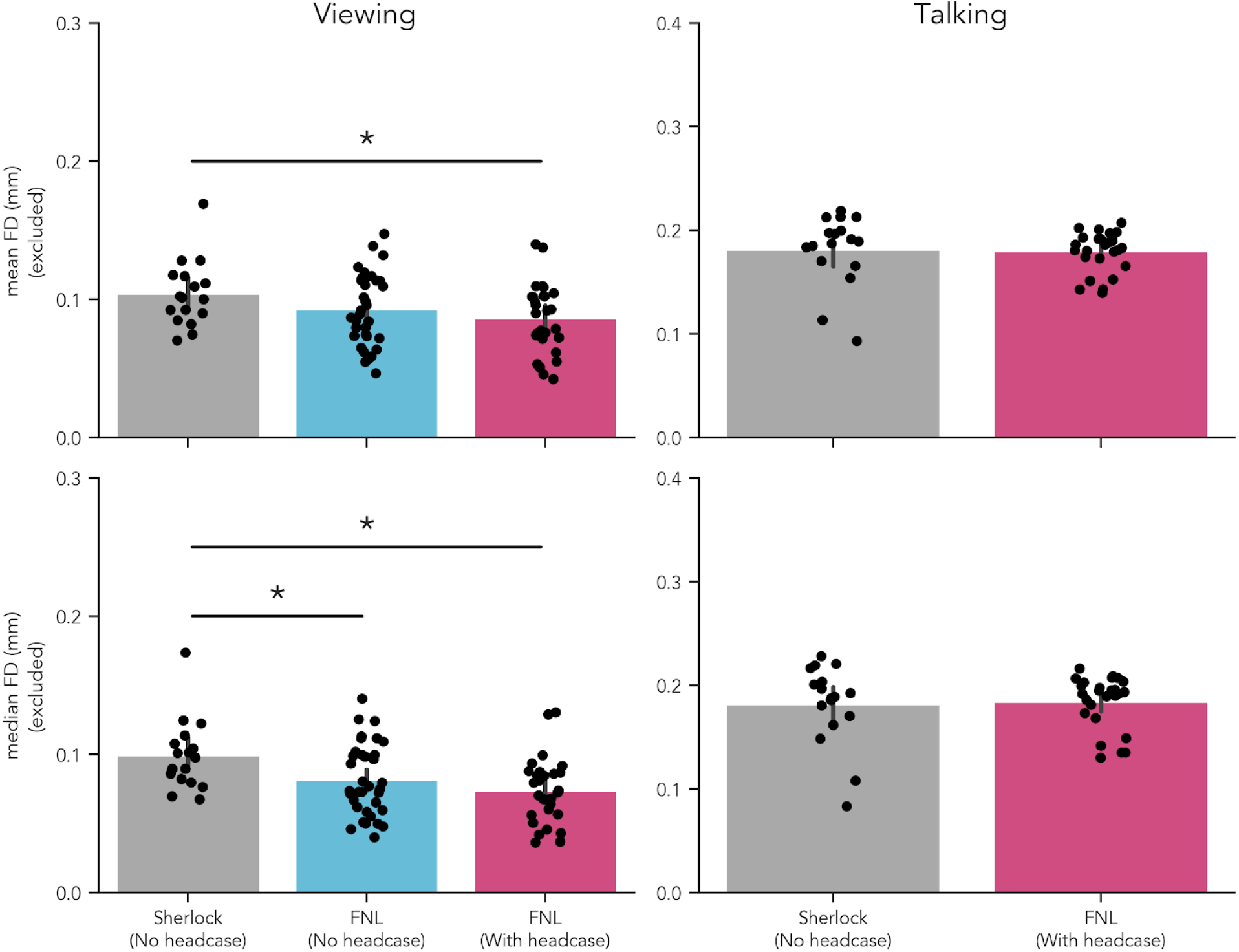
Headcase effects on Framewise Displacement excluding high motion volumes. Mean and median FD differences after removing high motion TRs (FD > 0.3mm). Headcases demonstrated no detectable improvement in small head motions either during viewing or during talking. However, the Sherlock sample showed increased median FD during viewing relative to both FNL samples. Motion in the Sherlock sample was significantly higher than both FNL samples during viewing. Mean FD was higher in the Sherlock sample relative to the FNL sample with headcases.

Our control analyses comparing FD_MeanExcluded_ and FD_MedianExcluded_ between our non-headcase sample and the *Sherlock* sample of participants without headcases produced similar results. We observed no significant difference in FD_MeanExcluded_, *t* = −1.541, *p* = .134, but did observe a difference in FD_MedianExcluded_ *t* = −2.368, *p* = .022. This translated to a significant FD_MeanExcluded_ difference between the *Sherlock* sample and *FNL headcase* sample *t* = 2.43, *p* = .022 as well as a significant FD_MedianExcluded_ *t* = 3.319, *p* = .002 (Fig 2 bottom row, left column; Table 3).

**Table 3.** Head case comparisons while *viewing* movies after excluding high motion TRs

| *Comparison* | *Condition* | *Metric* | $M_{\text{Difference}}$ | $t$ | $p_{\text{perm}}$ |
| --- | --- | --- | --- | --- | --- |
| FNL no-case - FNL with-case | Viewing | FD<sub>MeanFiltered</sub> | 0.008<br>(-0.005 0.021) | 1.186 | 0.244 |
| FNL no-case - FNL with-case | Viewing | FD<sub>MedianExcluded</sub> | 0.009<br>(-0.005 0.021) | 1.33 | 0.192 |
| FNL no-case - Sherlock | Viewing | FD<sub>MeanExcluded</sub> | -0.011<br>(-0.025 0.002) | -1.541 | 0.134 |
| FNL no-case - Sherlock | Viewing | FD<sub>MedianExcluded</sub> | -0.018<br>(-0.032<br>-0.004) | -2.368 | **0.022** |
| Sherlock - FNL with-case | Viewing | FD<sub>MeanExcluded</sub> | 0.019<br>(0.005 0.034) | 2.43 | **0.022** |
| Sherlock - FNL with-case | Viewing | FD<sub>MedianExcluded</sub> | 0.026<br>(0.012 0.042) | 3.319 | **0.002** |

### 3.3 Headcases result in more idiosyncratic motion across participants while viewing a movie

To test whether headcases impacted the degree to which participants exhibited common motion when viewing the same stimulus, we compared the average ISC of each pair of participants’ FD time-series in Datasets 1 and 2. The FNL with-case group (*M* = 0.02; *SD* = 0.04) exhibited significantly less shared motion compared to the FNL no-case group (*M* = 0.04; *SD* = 0.05), *t*(46.10) = 2.25, *p* = 0.029 (Fig 3, left panel). We also examined the ISC of each direction of motion separately to test whether headcases reduced common drifts in any direction reflected by each realignment parameter. None of these comparisons were significant, all *p*s > 0.40. However, the overall ISC of each motion direction in this analysis was very low and highly variable, making it difficult to detect any group differences (Fig S9).

**Figure 3.**
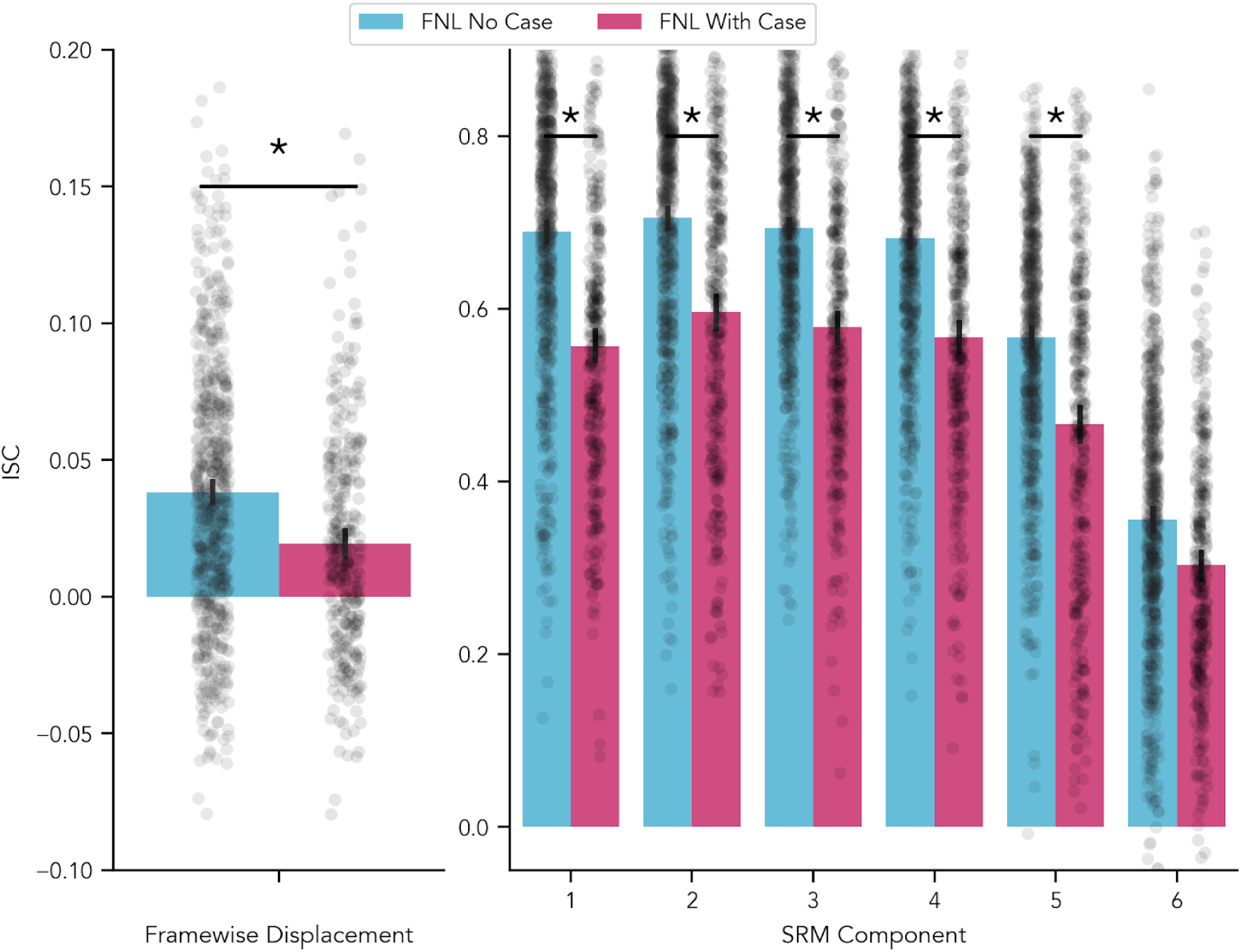
Headcase effects on shared motion. Intersubject correlation of Framewise Displacement (left) and aligned shared motion components (right). Participants with headcases (exhibited significantly less shared motion in their Framewise Displacement over time. By computing linear combinations of the six original realigned axes we were able to discover six new directions of motion (right) that maximally aligned across participants. In this new aligned space, participants with headcases exhibited significantly less shared motion in all but one component. This group difference was only apparent in one of the original unaligned motion parameters (Supplementary Materials; Fig S9).

To address this limitation, we relaxed the assumption that stimulus correlated movements must manifest in the same way for every participant. We instead computed a Shared Response Model (P.-H. (cameron) Chen et al., 2015) to discover a new set of latent motion axes that better capture shared motion across participants and repeated the group comparison for each shared component in this new space. Similar to the FD results above the FNL with-case group exhibited significantly less shared motion than the FNL no-case group across all components except one, p < 0.01 (Figure 3, right panel).

### 3.5 Lack of improvement in tSNR with headcases when viewing a movie

To examine whether headcases improved the overall quality of the measured fMRI signal we performed a comparison between the tSNR of each voxel for the FNL with-case and FNL no-case groups. Because both groups of participants watched the same stimulus collected with the same acquisition parameters, any statistical comparisons were unlikely to be due to differences in data collection procedures. Overall no voxels survived multiple comparisons correction, but a handful of voxels primarily on the dorsal and lateral edges of cortical gray matter showed a significant increase in tSNR at a more liberal p 0.001 uncorrected threshold (Figure 4). Mean group tSNR maps were highly similar for both groups suggesting that the spatial distribution of tSNR across gray matter voxels was highly consistent with or without headcases *r* = 0.984, *p* < .001. These results suggest that headcases had little measurable impact on the overall quality of the acquired signal when most other data acquisition procedures were matched between datasets.

**Figure 4.**
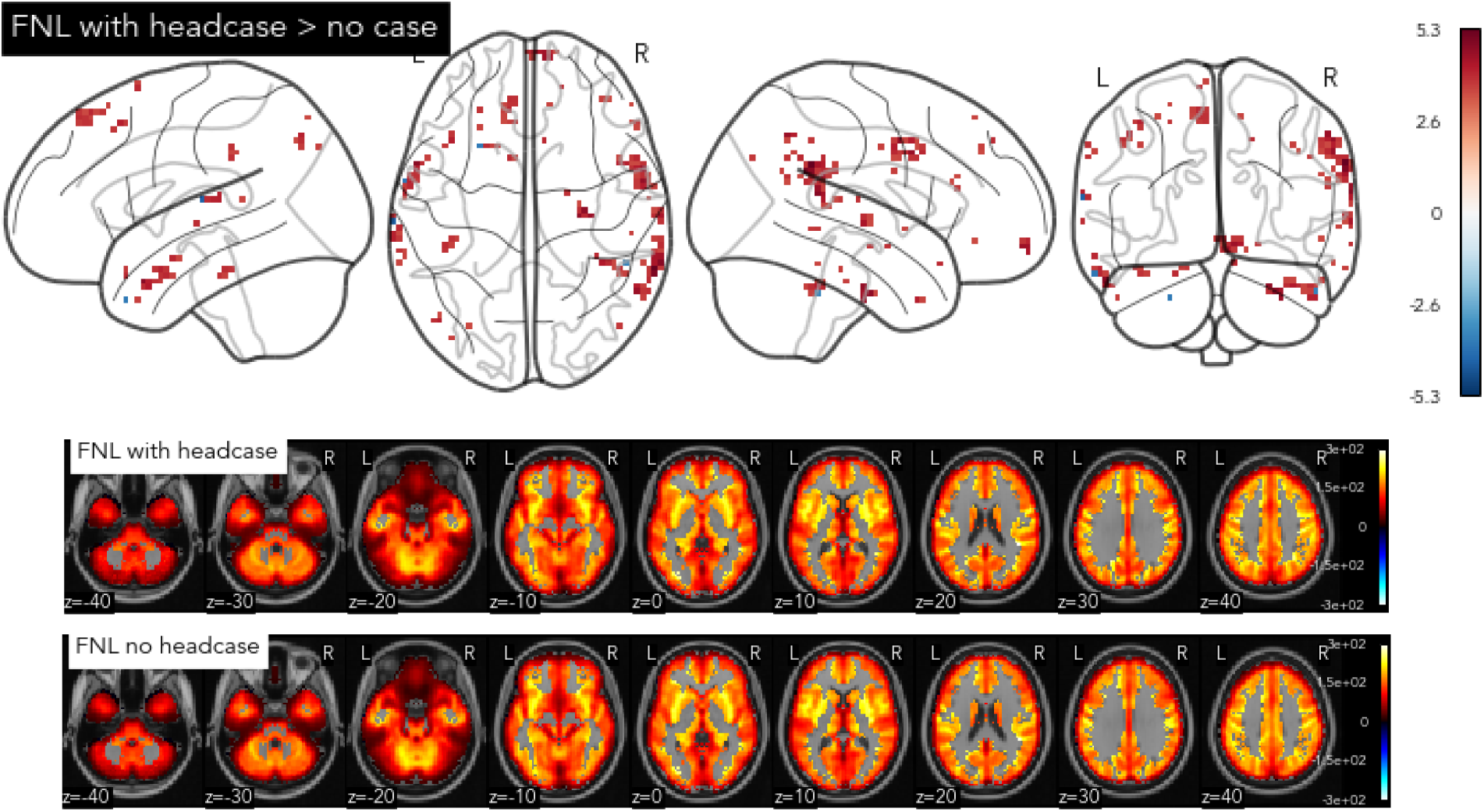
Headcase effects on tSNR. Group differences in voxel-wise tSNR between participants with and without headcases (top) and group mean tSNR maps (bottom). No voxels survived multiple comparisons correction when comparing tSNR with and without headcases. The contrast map above is thresholded at *p* < .001 uncorrected, in which few voxels primarily on the edges of the cortex show higher tSNR for participants wearing headcases relative to those not; values reflect t-statistic. Mean group tSNR maps (bottom; un-thresholded) were highly similar suggesting little improvement of signal quality with headcase use.

**Figure 5.**
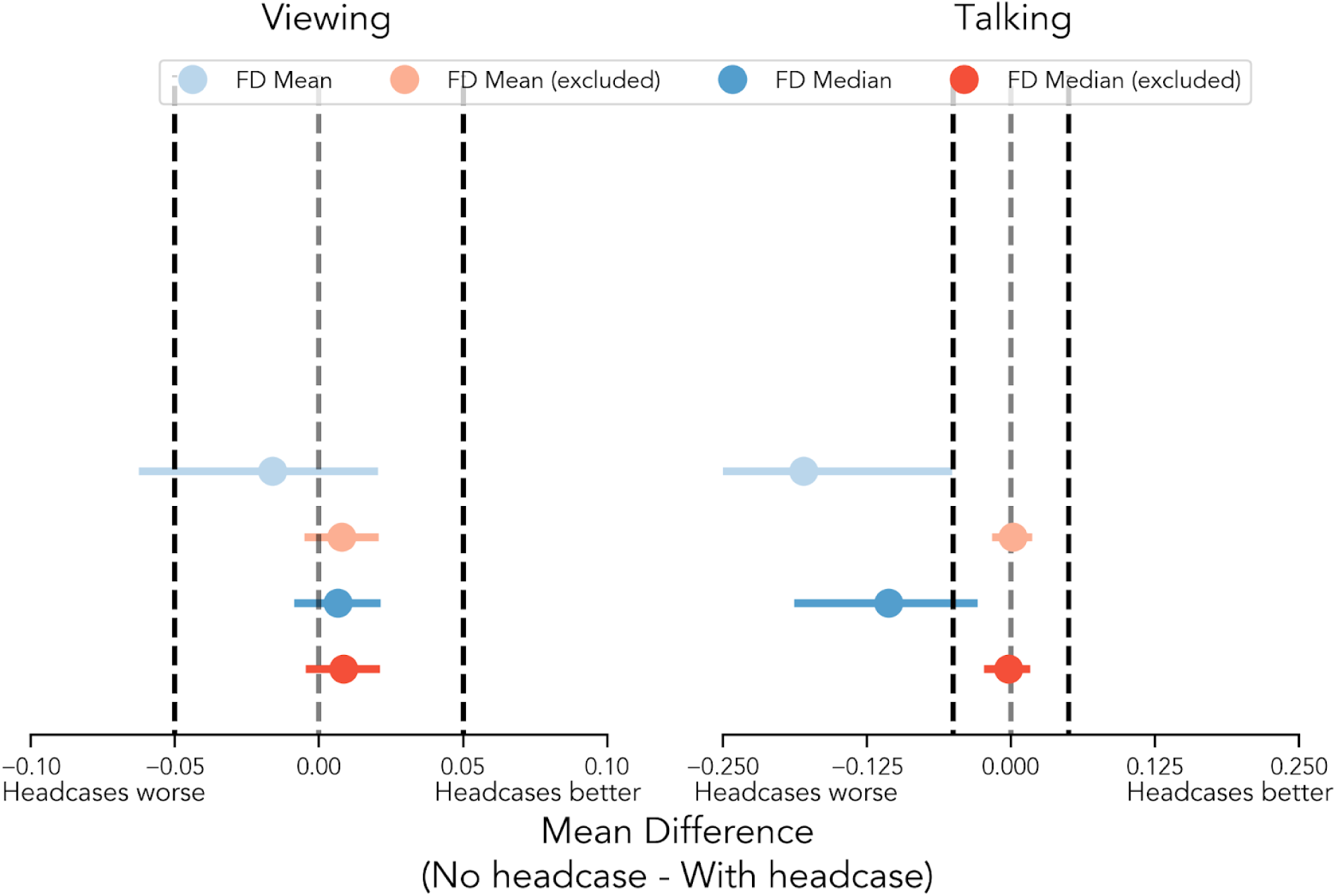
Equivalence tests of motion estimates with and without headcases. Plots depict whether mean differences between groups fall within bounds of practical value to fMRI researchers. Mean differences within these bounds can be interpreted as “statistically equivalent” (Lakens et al., 2018). Darker dashed lines indicate the upper and lower bounds of the equivalence tests, while the lighter dashed lines reflect a difference of 0. The upper positive bound (right dashed lines) in each plot tests whether head motion *decreased* by at least 0.05mm when wearing headcases. The lower negative bound (left dashed line) tests whether head motion *increased* by at least 0.05mm. In all cases, motion statistics were significantly below the upper equivalence bound suggesting that observed mean reductions in motion while wearing headcases are unlikely to be as large as 0.05mm. Mean differences that included high motion volumes during the talking condition (right blue points) also predominantly fell outside the lower equivalence bound, suggesting that in some situations headcases may exacerbate motion by at least 0.05mm.

### 3.6 Lack of overall motion reduction with headcases while talking aloud

While the previous analyses focused on a passive experimental “task” with a single long continuous run, our second set of analyses focused on an even more problematic scenario for motion in neuroimaging – speaking during the scan. While several recent studies have utilized verbalization tasks in the scanner (J. Chen et al., 2017; Silbert et al., 2014; Stephens et al., 2010; Zadbood et al., 2017), to our knowledge, none have compared the efficacy of custom headcases in mitigating talking-induced motion. We expected that even if headcases provided minimal reduction of head motion during passive viewing, perhaps due to increased participant compliance (Vanderwal et al., 2019), they *should* provide maximal benefit when participants move their heads as a consequence of task demands (i.e. talking). However, to our surprise, we did not find evidence supporting this hypothesis, and in some cases found the *opposite* result. When comparing Sherlock participants without headcases to our FNL with-case participants in the talking task, we found that participants wearing headcases exhibited *increased* motion across all metrics^2^ (Fig 1 right column). Participants wearing headcases exhibited higher FD_Mean_ (*M* = 0.467; *SD* = 0.362) compared to *Sherlock* participants (*M* = 0.287; *SD* = 0.116), *t* = −2.353, *p* = .009. They also exhibited higher FD_Median_ (*M* = 0.356; *SD* = 0.177) relative to participants without headcases (*M* = 0.25; *SD* = 0.097), *t* = −2.534, *p* = .013, as well as a larger proportion of high-motion volumes (Spike_Proportion_): with headcase (*M* = 0.516; *SD* = 0.237), without headcase (*M* = 0.345; *SD* = 0.23), *t* = −2.353, *p* = .025. These results suggest that headcases lack the efficacy to mitigate large head movements (e.g. > 0.3mm) observed in more problematic scenarios like speaking during a scan despite the extra restriction they place on participants (Table 4). No associations or differences between groups were observed between how long participants spoke for and average head motion (Supplementary Figures S4 & S5).

**Table 4.** Head case comparisons while *talking*. Excluded values reflect comparisons after excluding high motion TRs.

| *Comparison* | *Condition* | *Metric* | $M_{Difference}$ | $t$ | $p_{perm}$ |
| --- | --- | --- | --- | --- | --- |
| Sherlock - FNL with-case | Talking | FD<sub>Mean</sub> | -0.18<br>(-0.343 -0.051) | -2.353 | **0.009** |
| Sherlock - FNL with-case | Talking | FD<sub>Median</sub> | -0.106<br>(-0.188 -0.029) | -2.534 | **0.013** |
| Sherlock - FNL with-case | Talking | Spike<sub>Proportion</sub> | -0.171<br>(-0.306 -0.032) | -2.353 | **0.025** |
| Sherlock - FNL with-case | Talking | FD<sub>MeanExcluded</sub> | 0.002<br>(-0.016 0.018) | 0.190 | 0.857 |
| Sherlock - FNL with-case | Talking | FD<sub>MedianExcluded</sub> | -0.002<br>(-0.024 0.017) | -0.222 | 0.829 |

### 3.7 Lack of small motion reduction with headcases while talking aloud

We once again repeated the previous comparison after excluding high-motion volumes per individual to examine the efficacy of headcases for reducing smaller head movements while talking. Headcases demonstrated no significant reduction in FD_MeanExcluded_ between our sample (*M* = 0.18; *SD* = 0.019) and the *Sherlock* sample (M = 0.181; SD = 0.034) *t* = 0.19, *p* = .857. This was also true when examining FD_MedianExcluded_: with head case (*M* = 0.184; *SD* = 0.025), without headcase (*M* = 0.182; *SD* = 0.039), *t* = −0.222, *p* = .829. To further quantify these null results (Fig 5 right column), we again performed equivalence testing using the TOST procedure as described previously. We found that for both FD_MeanExcluded_ and FD_MedianFiletered_ observed differences in motion fell within this range of practical equivalence (Fig 5 right column, red points), *ps* < .001. These findings suggest that when specifically examining the reduction of smaller head movements induced while talking aloud, at best, headcases may be as efficacious as foam pillows alone (Table 4).

### 3.8 Causes of increased motion while wearing headcases and talking

Next, we explored what may have caused more motion while participants wearing headcases spoke aloud. We speculated that the interaction between lower jaw movements and head restriction may have paradoxically focused motion in the z-axis (moving head inward/outward parallel to the main axis of the scanner bore) and pitch-axis (nodding head up and down). In other words, we hypothesized that because participants were largely restricted in every direction but were unrestricted from moving their lower jaws, movements of the head may have been exacerbated in these planes. To test this hypothesis, we repeated our group comparison while participants talked *separately* for each axis of translation and rotation (Fig 6). Specifically, we compared displacement in the x, y, z, pitch, roll, and yaw axes using each participant’s mean, median, and standard deviation. Across all three summary statistics, we found significantly greater displacement in all rotation axes (pitch, roll, yaw), all *p*s < 0.05 (Table 5). We also found marginally greater mean displacement in the z-axis, *p* = 0.056, and significantly greater variability of displacement in both the x and z axes *p*s < 0.05. These findings suggest that observed differences in overall FD_Mean_, FD_Median_ and Spike_Proportion_ may have been driven by participants rotating their heads during talking, despite being constrained by headcases.

**Table 5.** Head case comparisons while *talking*. Values reflect mean differences and bootstrapped 95% confidence intervals for motion parameters without headcase – with headcase.

| *Motion Parameter* | *Metric* | *Mean<sub>Difference</sub>* | *t* | *p<sub>perm</sub>* |
| --- | --- | --- | --- | --- |
| X | Mean | -0.005<br>(-0.011 0.001) | -1.614 | 0.122 |
| Y | Mean | -0.01<br>(-0.037 0.009) | -0.838 | 0.588 |
| Z | Mean | -0.048<br>(-0.102 -0.005) | -1.898 | **0.056** |
| Pitch | Mean | -0.069<br>(-0.132 -0.02) | -2.356 | **0.007** |
| Roll | Mean | -0.031<br>(-0.045 -0.019) | -4.596 | **< 0.001** |
| Yaw | Mean | -0.016<br>(-0.028 -0.006) | -2.953 | **0.002** |
| X | Median | -0.001<br>(-0.005 0.002) | -0.608 | 0.552 |
| Y | Median | 0.001<br>(-0.009 0.01) | 0.171 | 0.874 |
| Z | Median | -0.017<br>(-0.043 0.004) | -1.425 | 0.166 |
| Pitch | Median | -0.031<br>(-0.055 -0.007) | -2.374 | **0.023** |
| Roll | Median | -0.018<br>(-0.025 -0.011) | -4.674 | **< 0.001** |
| Yaw | Median | -0.008<br>(-0.014 -0.002) | -2.748 | **0.011** |
| X | SD | -0.009<br>(-0.019 -0.001) | -1.890 | **0.043** |
| Y | SD | -0.026<br>(-0.075 0.004) | -1.141 | 0.216 |
| Z | SD | -0.082<br>(-0.179 -0.01) | -1.838 | **0.040** |
| Pitch | SD | -0.105<br>(-0.231 -0.026) | -1.896 | **0.004** |
| Roll | SD | -0.038<br>(-0.061 -0.021) | -3.574 | **0.000** |
| Yaw | SD | -0.027<br>(-0.057 -0.008) | -2.009 | **0.001** |

**Figure 6.**
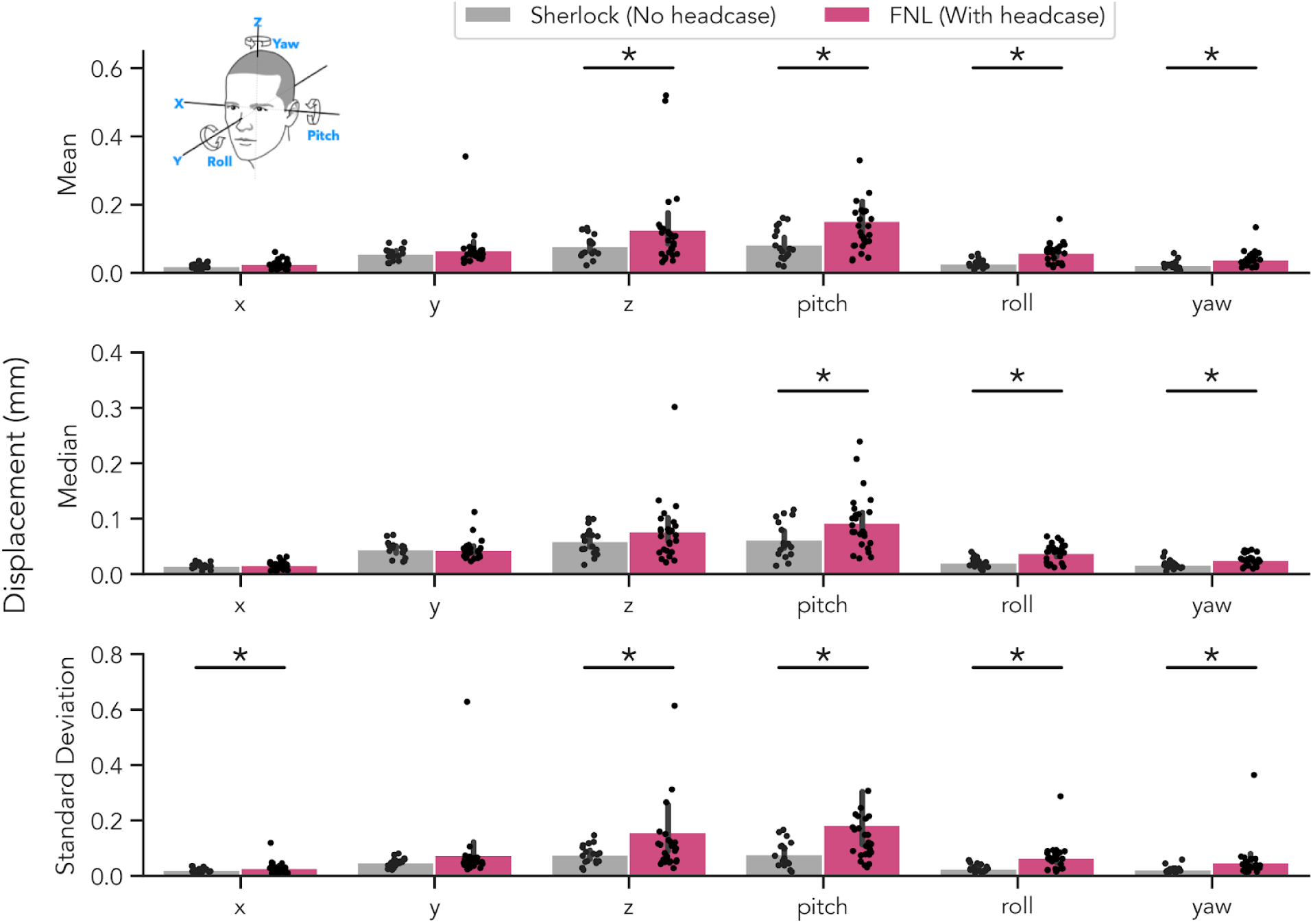
Headcase effect on individual motion parameters while talking. Differences in average displacement for each motion parameter as a function of headcase use while talking (Sherlock and FNL samples). Each plot depicts the absolute value of the backwards-differenced parameter estimate (i.e. the values ultimately summed to computed FD). Top row: average of each participant’s mean displacement; middle row: median displacement; bottom row: standard deviation of displacement. In all metrics, headcase participants demonstrated significantly more displacement in all rotation directions (pitch, roll, yaw) with pitch demonstrating the largest differences, likely as a result of bottom-jaw induced head movement. Significant differences were also observed in the z-axis for mean displacement and the x and z-axes for standard deviation of displacement.

## 4. Discussion

Using three fMRI datasets, we tested the efficacy of custom-molded headcases in reducing head motion during longer naturalistic tasks (viewing a movie or talking aloud). Unlike previous work (Jonathan D. Power et al., 2019), which demonstrated an overall reduction in head motion during a brief resting state scan, we found that headcases provided little benefit in reducing head motion. Between group comparisons (with and without headcase) of movie-watching indicated no significant reduction in overall head motion indexed by mean and median Framewise Displacement (FD), nor reduction in the proportion of high-motion volumes sampling idiosyncrasies as participants were recruited from the same population (Dartmouth College community) and scanned using the same equipment with identical acquisition parameters and stimulus. On the contrary, this population as a whole displayed less overall motion relative to a similar population that also underwent a naturalistic movie-viewing paradigm collected at a different site (Princeton University) (J. Chen et al., 2017). Additionally, overall motion estimates for participants in our samples were approximately similar to participants in the same age range within the sample examined by Power et al (2019).

Interestingly, we did find that participants who wore headcases displayed more idiosyncratic motion that those who did not. This was reflected by a higher ISC of the FD time-series for non-headcase participants. While FD captures momentary displacements based on successive differences in head position, we also found that participants without headcases displayed more correlated drifts in motion. Specifically, we estimated unique transformation matrices that separately projected each participants’ original motion parameters into a new latent shared motion space. In this new space, participants without headcases had higher ISC across nearly all shared components. The high overall magnitude of the ISC for both groups in this shared motion space was particularly notable. Though our ISC estimate is likely to be slightly inflated due to training and testing the model on the same data, it is unlikely to induce group differences where none exist, especially given the ISC of FD time-series we observed. One potential interpretation of these findings is that headcases are effective at reducing stimulus correlated motion (i.e. shared motion induced by watching a movie) despite not decreasing overall head motion and that this shared motion is not captured well by displacements and rotations about the axes captured by the original realignment parameters (Fig S9). Common motion, potentially induced by the stimulus, appears to be highly preserved across individuals despite a significant reduction from headcase use. Determining whether this shared motion is caused by the stimulus itself presents an interesting possibility for future research, but will require many additional analyses that are beyond the scope of the current paper. These findings suggest that accounting for motion during preprocessing may be a critical analysis step for naturalistic studies.

Surprisingly, we found that participants who wore headcases produced *more* motion than those who did not when speaking aloud inside the scanner. This seems to have been driven by individual volumes with large movements (FD exceeding 0.3mm), as group comparisons excluding these volumes yielded no significant differences between groups. These larger movements occurred in specific translation directions (i.e., z-axis) and across all rotational axes during talking. These findings suggest that headcases may be inadequate at restricting larger movements and in fact, may paradoxically amplify motion in scenarios that researchers might expect head motion to be worse (talking). With this finding in mind, we speculate that it is possible that by trying to restrict head motion while moving their lower jaws, participants rotate their heads *more* when wearing headcases (relative to foam pillows) increasing overall motion.

These null differences along with equivalence tests (Lakens et al., 2018) using effect sizes from previous work (Jonathan D. Power et al., 2019) highlight the limited efficacy of headcases in reducing head motion. Researchers must balance a tradeoff between the added time, money, and effort of data quality improvement procedures and their expected benefit. For Caseforge headcases specifically, researchers must order a special 3D camera from the company, photograph each participant in a separate session prior to MRI scanning, and await the arrival of each case. In some cases, this procedure must be repeated if cases contain defects or fit poorly. In our experience, nearly 20% of participants experienced discomfort when using headcases for extended periods of time precluding their use altogether or requiring adjusted procedures like using the front or back of the case only. This adds additional time and logistical challenges to data collection on top of the existing challenges that collecting MRI data already requires. Because headcases are by definition personalized for each individual, they offer no reusability if these same participants are no longer available for future scanning sessions. This may encourage researchers at a given institution to repeatedly sample a small subset of participants for whom headcases exist, decreasing the generalizability of empirical findings (Henrich et al., 2010; Yarkoni, 2019).

In the datasets analyzed here, each participant without a headcase was situated with foam pillows only (Dataset 3) or both foam pillows and medical tape (Dataset 1). We found this procedure to be adequate and flexible in a non-clinical, non-developmental population without the added burden of acquiring headcases. Interestingly, we found that participants for whom medical tape was used actually produced less overall motion relative to those who were situated with foam pillows alone (FNL no headcase vs Sherlock). We believe that this was driven by the tactile feedback on participants’ foreheads provided by the tape rather than any additional movement constraints. This is consistent with recent findings by Kraus et al (2019) who observed less within-run, between-run, and drifting head motion when using medical tape. Specifically, they observed that the efficacy of tactile feedback scaled with the amount of motion observed without tactile feedback, making this procedure particularly appealing when participants are likely to move a lot. Their findings also appear to be independent of behavioral tasks performed in the scanner and are efficacious even in situations when a participant is deliberately asked to move their head (Krause et al., 2019). Another promising alternative is the use of inflatable head pillows (e.g. Pearltec MULTIPAD Positioning System, Newmatic Medical, 2020) which may offer a compromise between completely head conforming designs like headcases and more general approaches like foam pillows. However, to our knowledge a systematic investigation using inflatable pillows has not yet been conducted.

The goal of the present work is to provide an in-depth analysis of the utility of headcases in reducing head motion^3^. Currently, there is only a single published investigation on the efficacy of headcases conducted using a small sample of 13 individuals and a short rsfMRI scan of 4.8 minutes (Jonathan D. Power et al., 2019). Our work adds to the existing literature to help researchers make a more informed decision about their data collection procedures. We are grateful that companies like Caseforge are working on developing fast, customizable, and accessible pipelines to battle the problematic issue of head motion for fMRI and hope that these results may lead to improvements in the design and manufacturing processes. Through private communication with Caseforge (Gao, 2019), we were notified that the headcases used in our sample (and presumably those used by Power et al (2019)) were the first generation of their kind and that more improvements to the reliability and comfort of headcases have been made in the second and forthcoming generations. While we are unable to test these claims, they provide promise for reducing the attrition rate we observed and further reducing head motion. Our sample also consisted of a non-clinical, non-development population. Such populations often exhibit increased head motion, a scenario in which headcases may provide more observable benefits (Vanderwal et al., 2015). The figures presented in Power et al (2019) support this notion as the largest motion reductions occurred for younger participants (7-14 years old). We speculate as to whether training procedures combined with tactile feedback might be similarly successful as participants can be taught to monitor their own movements (Krause et al., 2019).

In conclusion, we provide data and comparisons investigating the effect of headcases on reducing head motion under highly demanding data acquisition conditions: long task-free runs of movie-watching and active verbalization. Unlike previous work, we find that customized headcases provide no significant benefits for the reduction of head motion in a non-clinical, non-developmental population. We encourage future researchers to perform additional comparisons using alternative procedures for reducing headmotion (e.g., inflatable pillows, medical tape, headcases, etc). Together these investigations can better help the broader neuroimaging community adopt the most ideal practices to mitigate motion from contaminating fMRI data (Zaitsev et al., 2015).

## Open Practices Statement

All code and data required to reproduce the analyses and figures in this manuscript is available on github at https://github.com/cosanlab/headcase and on OSF at https://osf.io/qf6vx. A preprint of this manuscript is available on bioRxiv.

## Acknowledgements

The authors wish to thank Janice Chen & Uri Hasson for sharing their data and helpful comments on drafts on this manuscript, as well as James Gao and the Caseforge team for assistance and guidance in 3D photography used for the purchase of headcases as well as for helpful comments on drafts of this manuscript.

## Author contributions

E.J. and S.S. collected and analyzed the data. E.J., S.S., and L.J.C. wrote the manuscript.

## Footnotes

1 Reporting conventions for TOST results typically reflect the *higher* p-value of each one-sided test, except in cases where the one-sided test against a specific bound of equivalence is of primary interest. The reported p-values for all equivalence tests are for the *upper bound* of 0.05mm such that headcases yield a motion *decrease* of at least 0.05mm which is likely of more interest to fMRI researchers.

2 We reran these comparisons excluding two headcase participants who exhibited particularly high-levels of motion overall (FD_Mean_ >= 1mm; data points not depicted in Figures), but found that headcase participants still exhibited significantly higher FD_mean_ albeit equivalent levels of motion reflected in FD_Median_ and Spike_Proportion_ (see Supplementary Fig S1 & S2).

3 Head cases have also been advertised to ensure that an individual’s head is in the same position at the iso-center of the magnetic and head coil across sessions. We were unable to evaluate this in the present work and instead focused on motion mitigation.

