## Supplementary Materials for "Custom-molded headcases have limited efficacy in reducing head motion during naturalistic fMRI experiments"

**Figure S1**
Framewise Difference comparisons excluding 2 high motion participants in the FNL with headcase dataset.

**Figure S2**
Framewise Difference comparisons excluding 2 high motion participants in the FNL with headcase dataset and excluding high motion TRs

**Figure S3**
Equivalence test comparisons excluding 2 high motion participants in the FNL with headcase dataset.

**Figure S4**
Comparison between run length during *talking* and average Framewise Displacement.

A permuted regression analysis with bootstrapped confidence intervals indicated no significant relationship between run length and $FD_{Mean}$ $b$ = -0.0003 [-.00008 .0001], $t$ = -0.986, $p$ = 0.331. No interaction was observed between wearing headcases and run length $b$ = 0.0003 [ -.0001 .0008], $t$ = 0.813, $p$ = 0.440.

**Figure S5**
Comparison between run length during *talking* and average Framewise Displacement excluding 2 high motion participants in the FNL with headcase dataset.

A permuted regression analysis with bootstrapped confidence intervals indicated no significant relationship between run length and $FD_{Mean}$ $b$ = -0.0001 [-.00004 .0002], $t$ = -0.739, $p$ = 0.470. No interaction was observed between wearing headcases and run length $b$ = 0.0001 [ -.0002 .0005], $t$ = 0.617, $p$ = 0.537.

**Figure S6**
Mean difference in Framewise Displacement over time (FNL *viewing*).

Cumulative mean differences in FD were computed between the FNL with-case and FNL no-case groups using increasing window sizes of 30s (e.g. 30, 90, 120, etc). The shaded region reflects bootstrapped 95% confidence intervals of each mean difference estimated by resampling participants from each group with replacement 5000 times while preserving group sizes. The dashed black line reflects a difference of 0 between groups. Consistent with comparisons of $FD_{Mean}$, participants with headcases moved more than participants without for most of the scan duration. While the mean difference between groups appears to favor headcase early in the scan (i.e. < 4 minutes) confidence intervals include 0 suggesting that this difference is not statistically significant.

**Figure S7**
Individual linear regression estimates (black) for each participant's FD time-series (FNL *viewing*).

To examine whether FD exhibits a linear drift over the run and whether this drift differs with headcase use, we fit a multilevel model predicting FD using a linear Legendre polynomial, an indicator variable for headcase, and an interaction term between them with random intercepts and slopes per participant. No predictors were significant: linear trend $b$ = 0.01, $t$(61) = 1.85, $p$ = 0.069; headcase use $b$ = -0.02, $t$(61.01) = -0.865, $p$ = 0.395; interaction $b$ = 0.002, $t$(61) = 0.19, $p$ = 0.850. These results suggest that FD does not exhibit a reliable linear drift across participants and headcases provide no measurable impact on the linear drift of FD.

**Figure S8**
Individual logistic regression estimates (black) for each participant's spike probability (TR > 0.3mm; FNL *viewing*).

To examine whether spike frequency exhibits a linear drift and whether this drift interacts with headcase use we fit a logistic multilevel model predicting the probability of a spike (i.e. TR with > 0.3mm FD) using a linear Legendre polynomial, an indicator variable for headcase, and an interaction term between them with random intercepts and slopes per participant. We found a strong significant relationship between the linear term and spike probability $b$ = 0.442, $z$ = 3.143, $p$ = 0.002; no effect of headcase use $b$ = -0.056, $z$ = -0.140, $p$ = 0.889; and no interaction $b$ = 0.060, $z$ = 0.344, $p$ = 0.731. This suggests that the likelihood of large brief movements increase with time, but headcases do not mitigate this likelihood.

**Figure S9**
Group differences in ISC of each motion parameter between participants who did and didn't wear headcases (FNL *viewing*).

**Figure S10**
Time-frequency plots for each dataset visualizing detectable stable frequencies in motion parameters.

Following the approach in Fair et al (2020) the power spectra of each participant's six realignment parameters was computed using multi-taper spectral density estimation. Because the sampling frequency was limited by the TR in each dataset (Datasets 1 & 2: TR = 2s; 0.5hz; Dataset 3: TR = 1.5s; 0.66hz), reliable power estimation was limited to maximum of 0.25hz and 0.33hz without aliasing as given by the nyquist theorem. Each plot reflects relative power in each frequency band scaled between 0-100 using minmax normalization for comparability. Rows in each figure reflect individual participants and are sorted by $FD_{Mean}$ with low movers at the top and high movers at the bottom. As expected, participants with higher $FD_{Mean}$ had more detectable broadband power across several motion parameters. At the same time, we also observe increased narrow band power at low frequencies in both talking datasets, consistent with motion of the head as individuals speak. However, unlike previous work no identifiable respiration related artifact was detectable in any datasets. While the source of the inconsistency with previous work is unclear, one possibility is the lack of high temporal resolution afforded by multi-band acquisition which were not employed in any of the datasets here.

**Table 1**
Framewise Difference statistics during *viewing* excluding 2 high motion participants in the FNL with headcase dataset

**Table 2**
Framewise Difference statistics during *viewing* excluding high motion volumes and 2 high motion participants in the FNL with headcase dataset

**Table 3**
Framewise Difference statistics during *talking* (both with and without high motion volume) excluding 2 high motion participants in the FNL with headcase dataset

**Figure S1. Headcase effects on Framewise Displacement when viewing and speaking inside the scanner, excluding 2 high motion participants from the FNL with headcase dataset**

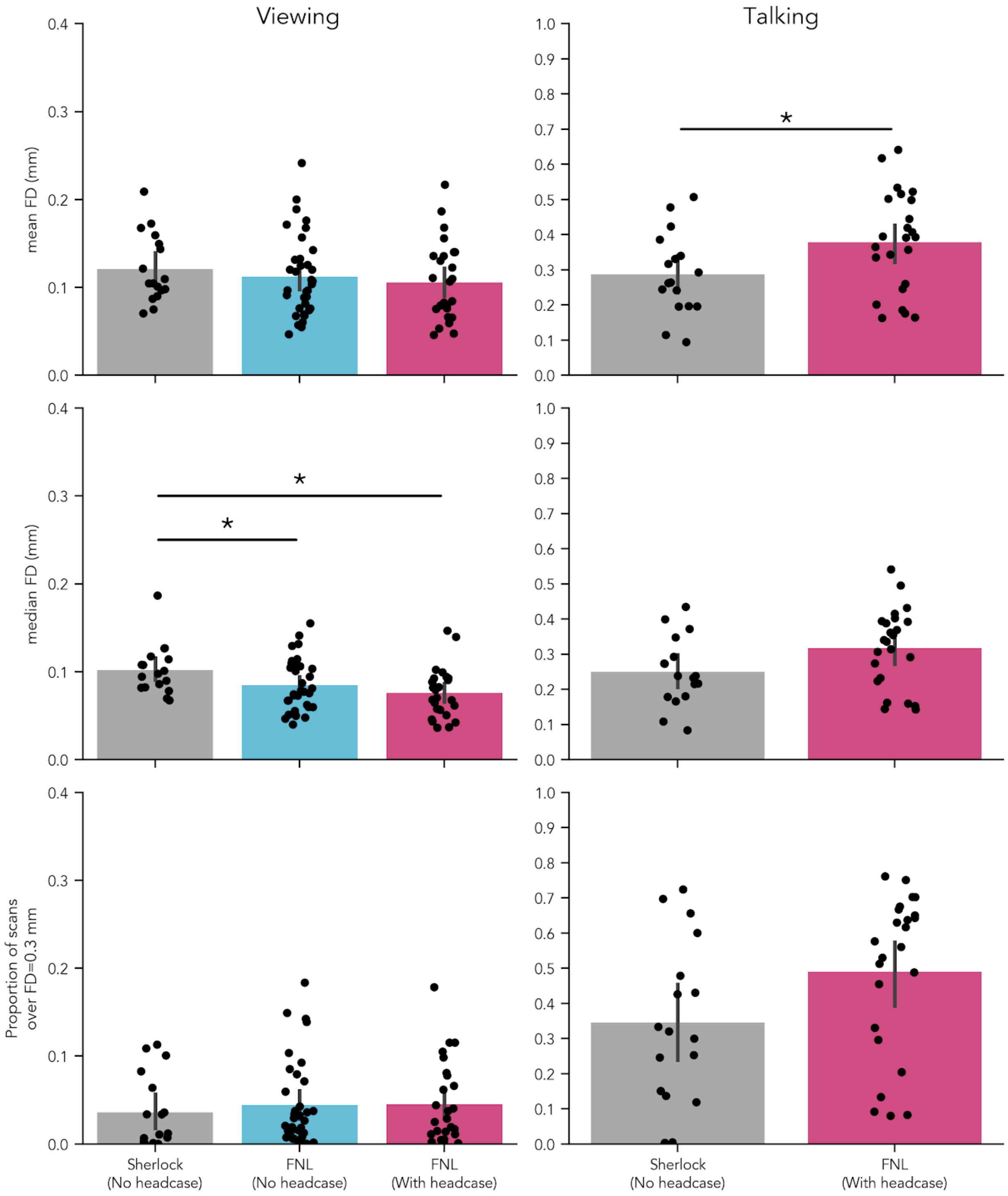

**Figure S2. Headcase effects on Framewise Displacement excluding high motion volumes and excluding 2 high motion participants from the FNL with headcase dataset**

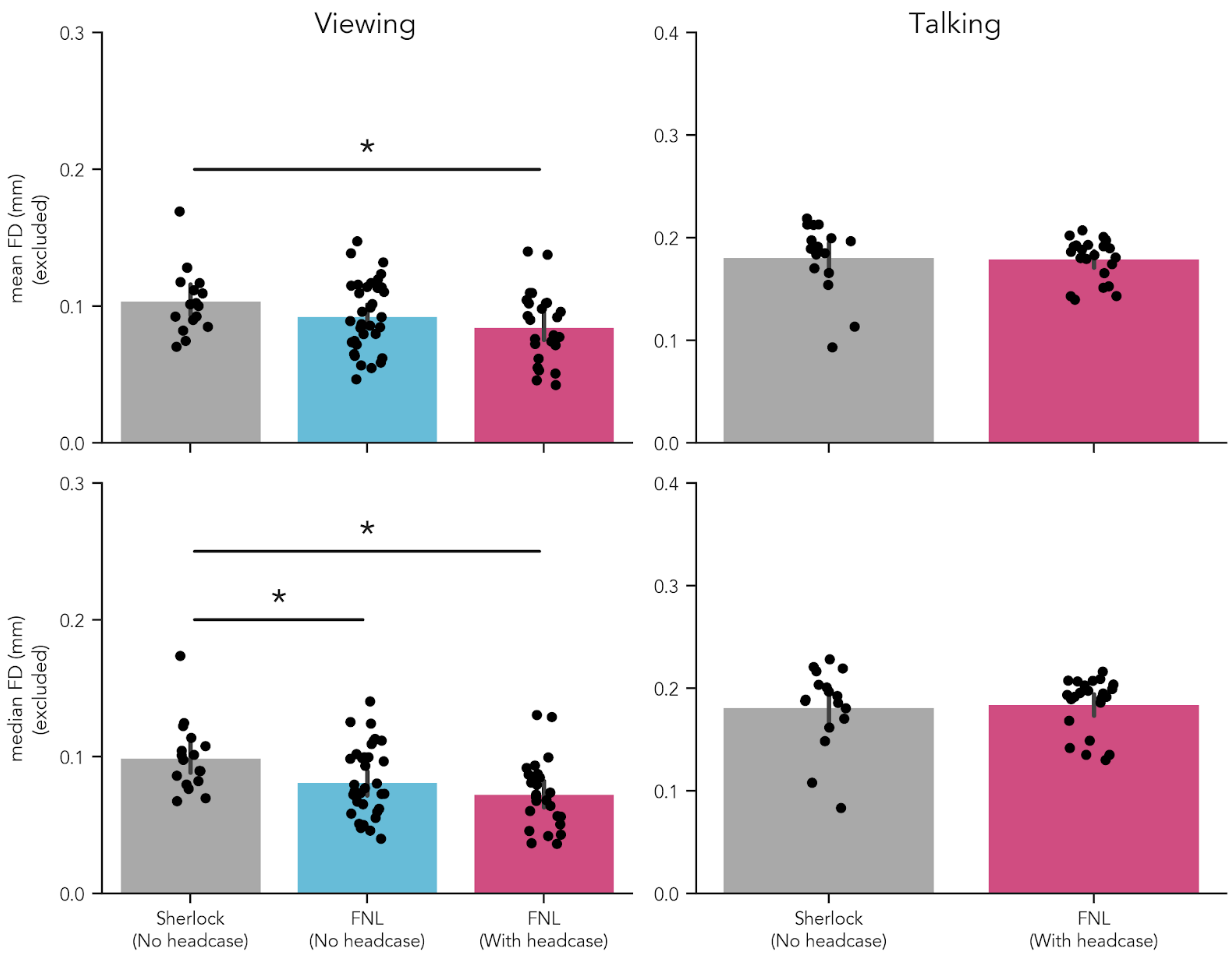

**Figure S3. Equivalence tests of motion estimates with and without headcases, excluding 2 high motion participants from the FNL with headcase dataset**

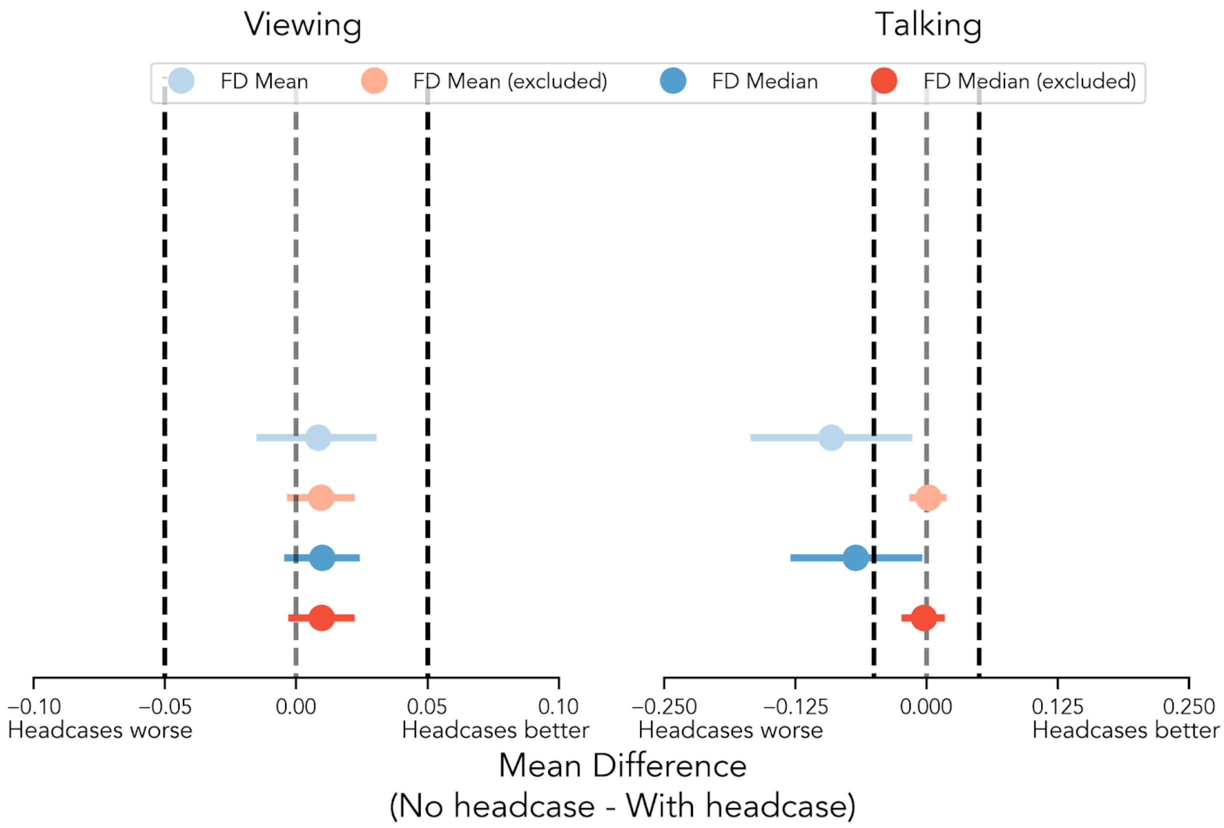

**Figure S4. Comparison of relationship between recall length and average head motion between Sherlock and FNL with headcase datasets.**

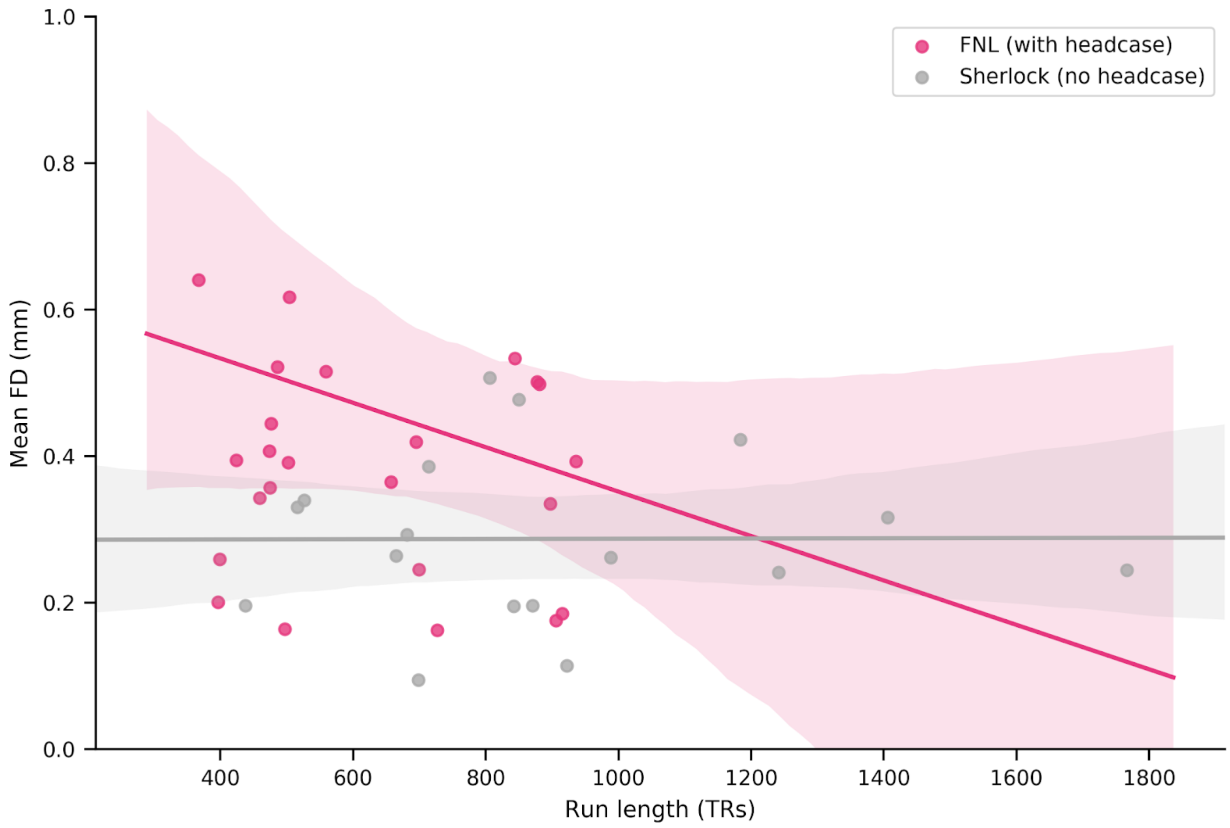

**Figure S5. Comparison of relationship between recall length and average head motion between Sherlock and FNL with headcase datasets, excluding 2 high motion participants in FNL with headcase dataset.**

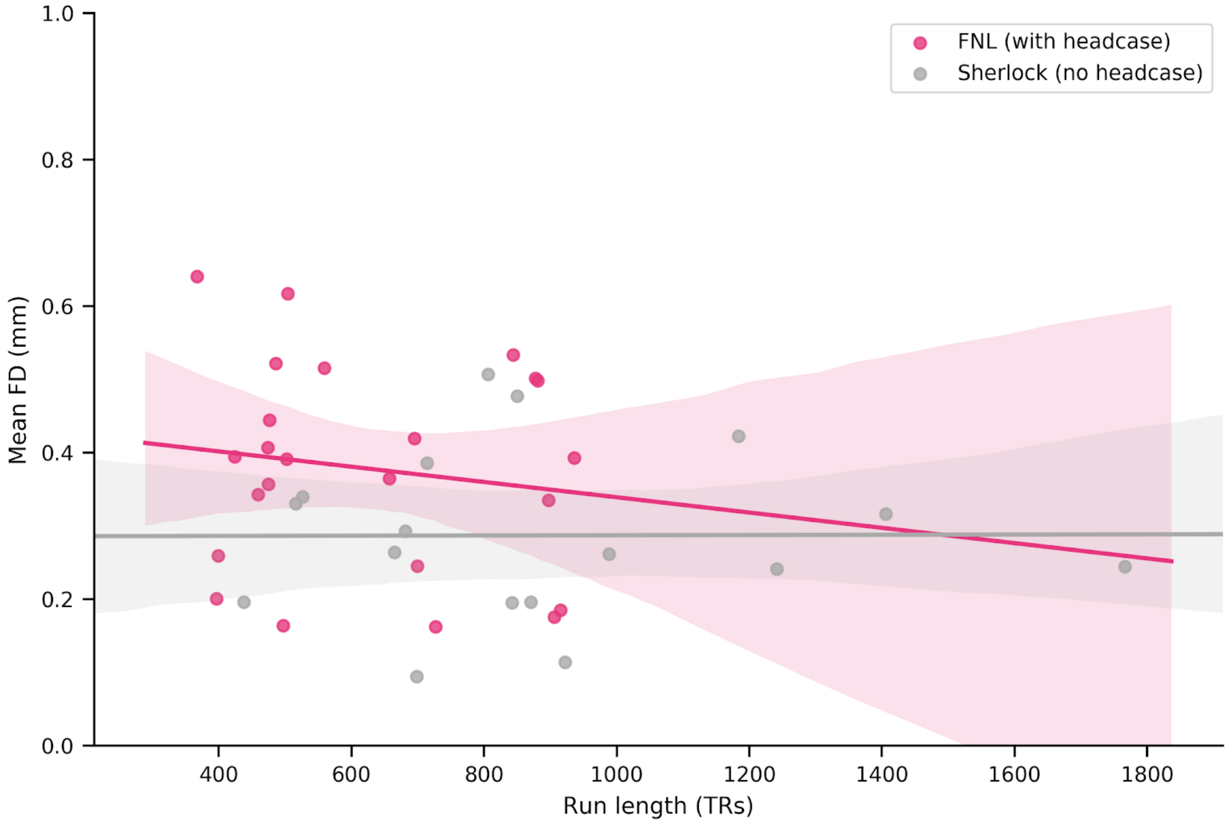

**Figure S6. Mean difference in Framewise Displacement over time while *viewing* FNL**

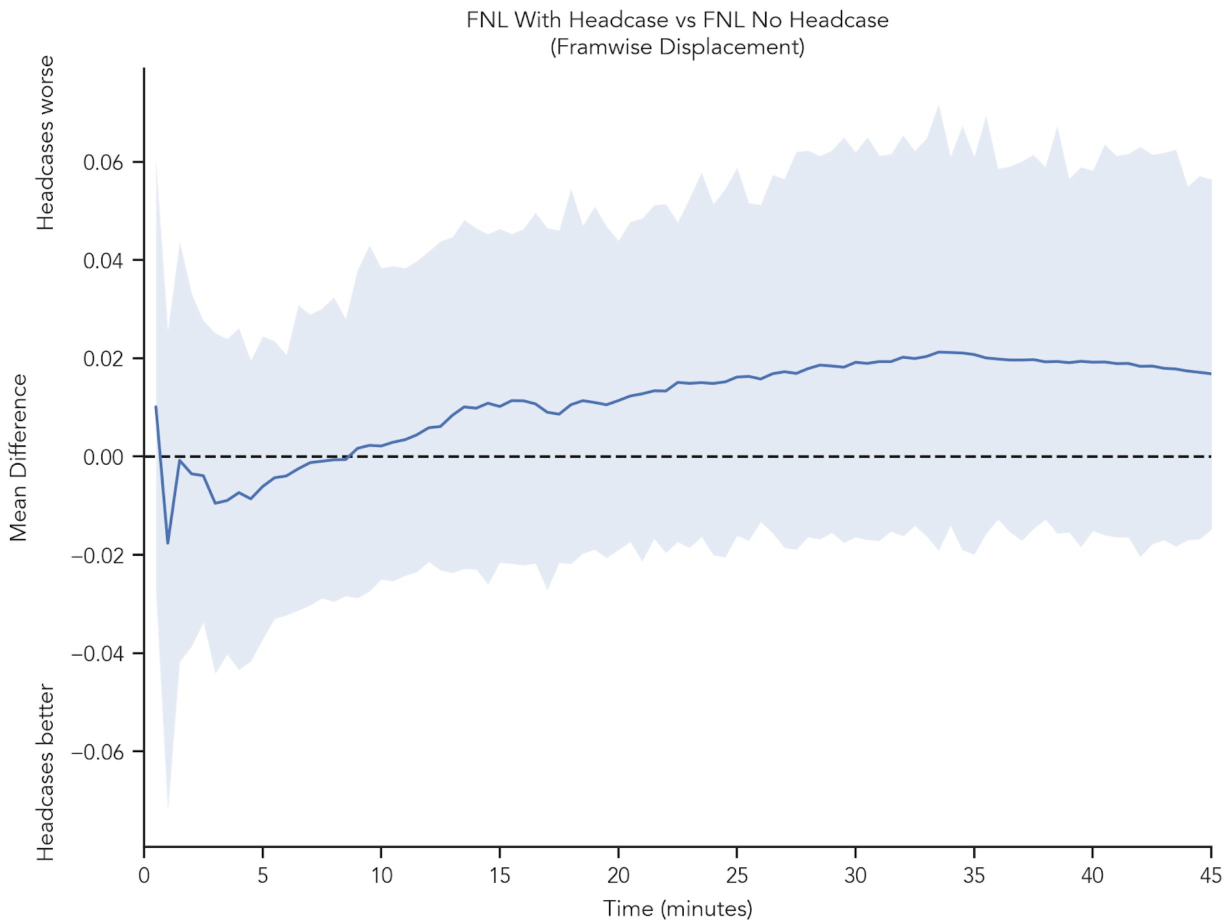

**Figure S7. Individual participant linear drift estimates of Framewise Displacement while *viewing* FNL**

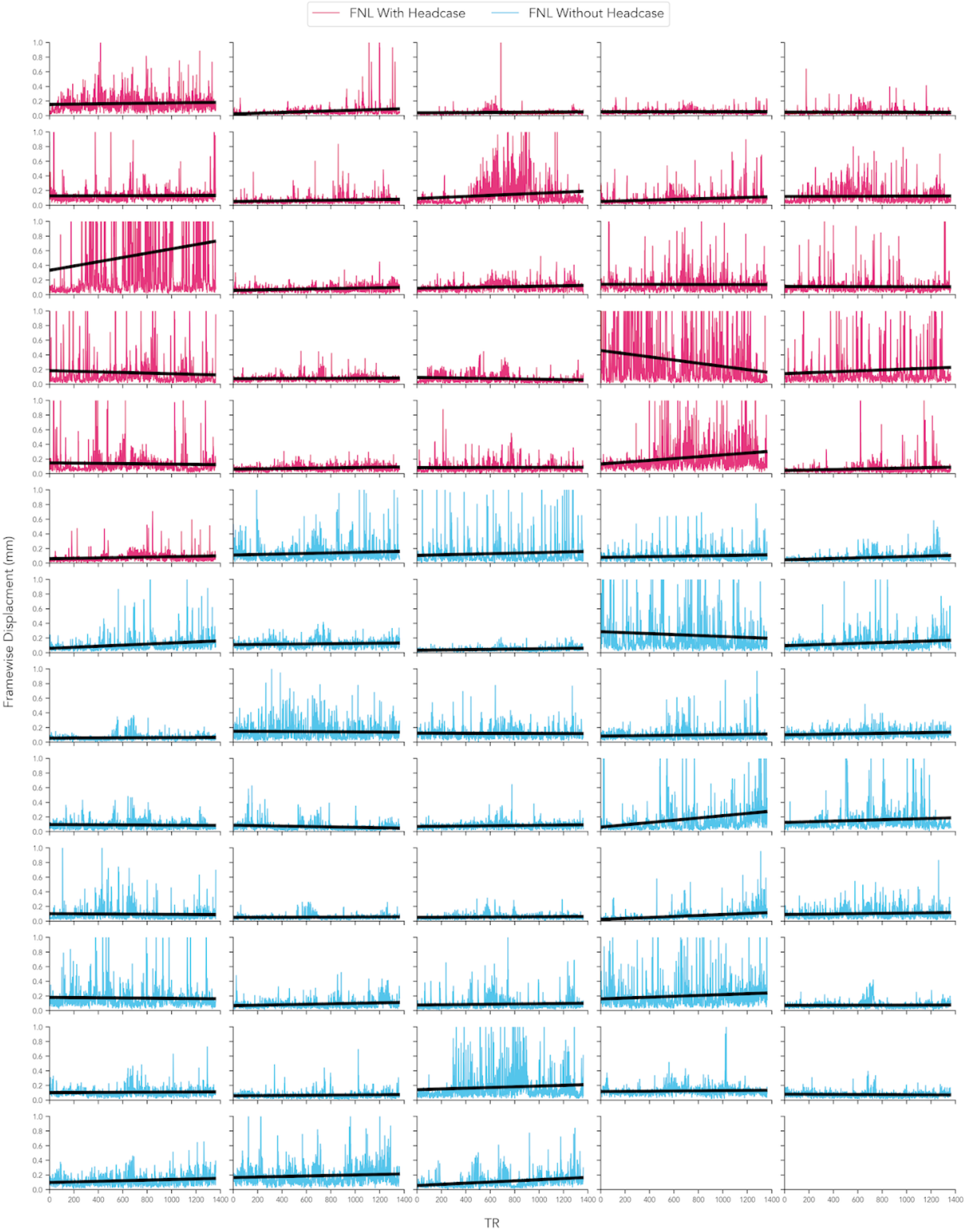

**Figure S8. Individual participant logistic motion spike likelihood estimates while *viewing* FNL**

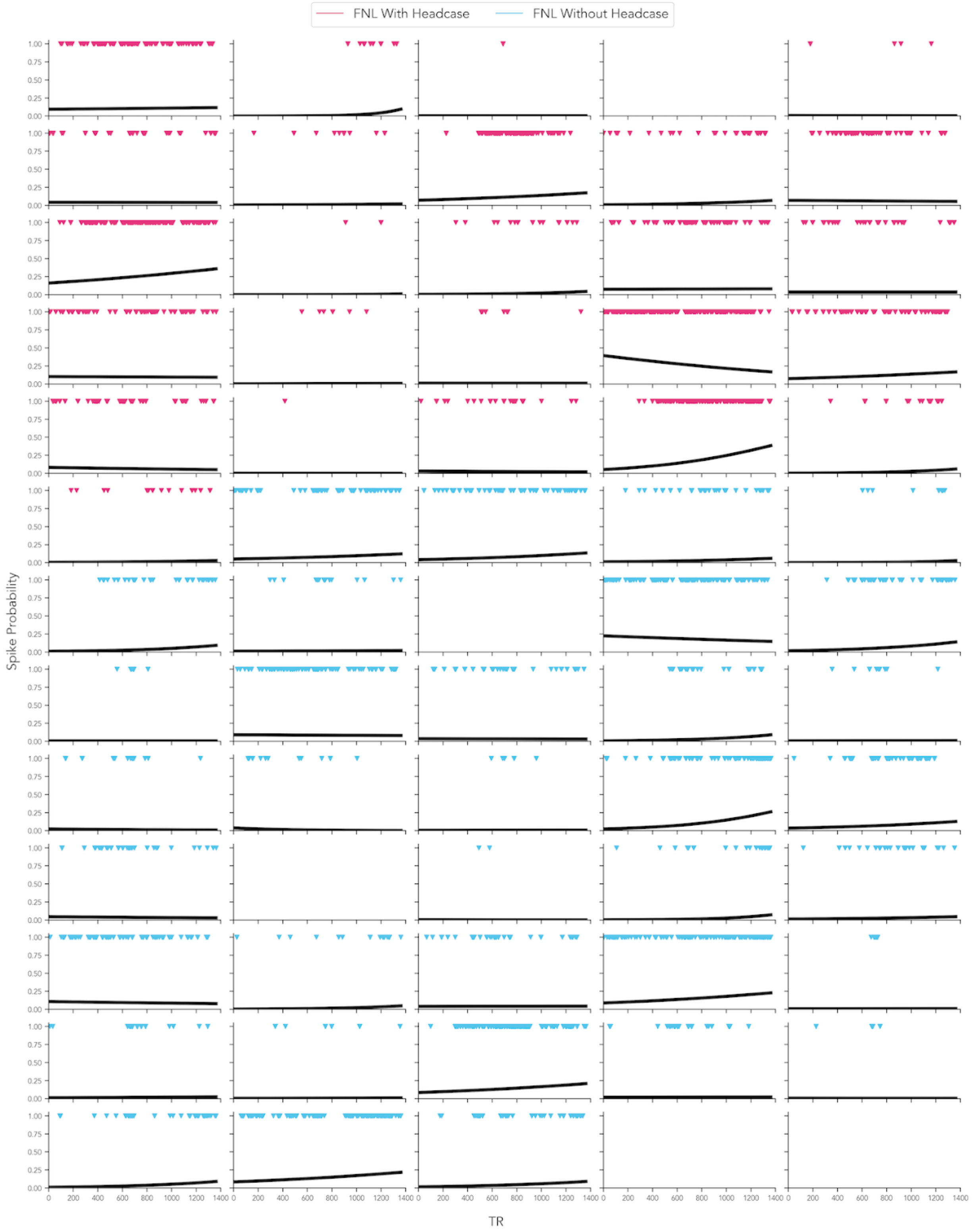

**Figure S9. Group differences in ISC of each motion parameter between participants who did and didn't wear headcases (FNL *viewing*).**

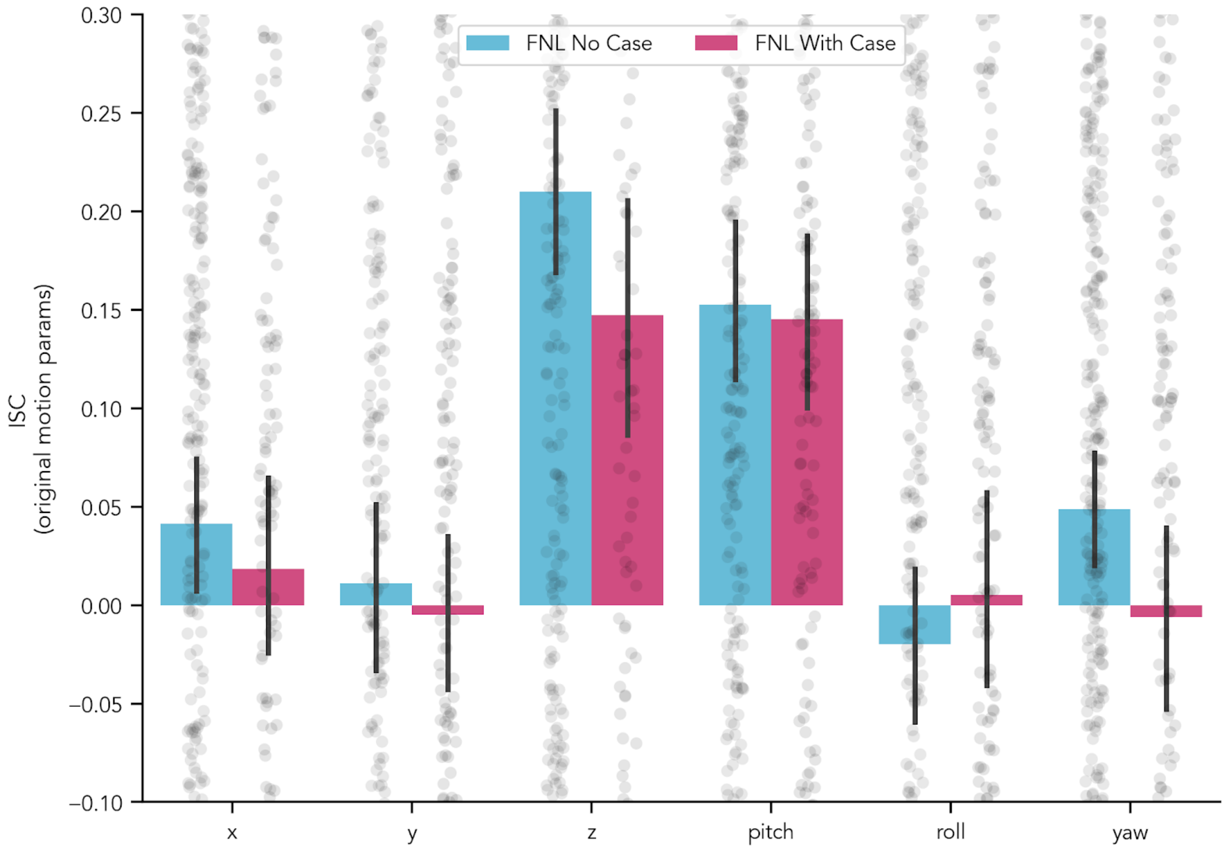

### S10. Time-frequency power spectra of motion parameters estimated via multi-taper spectral density estimation

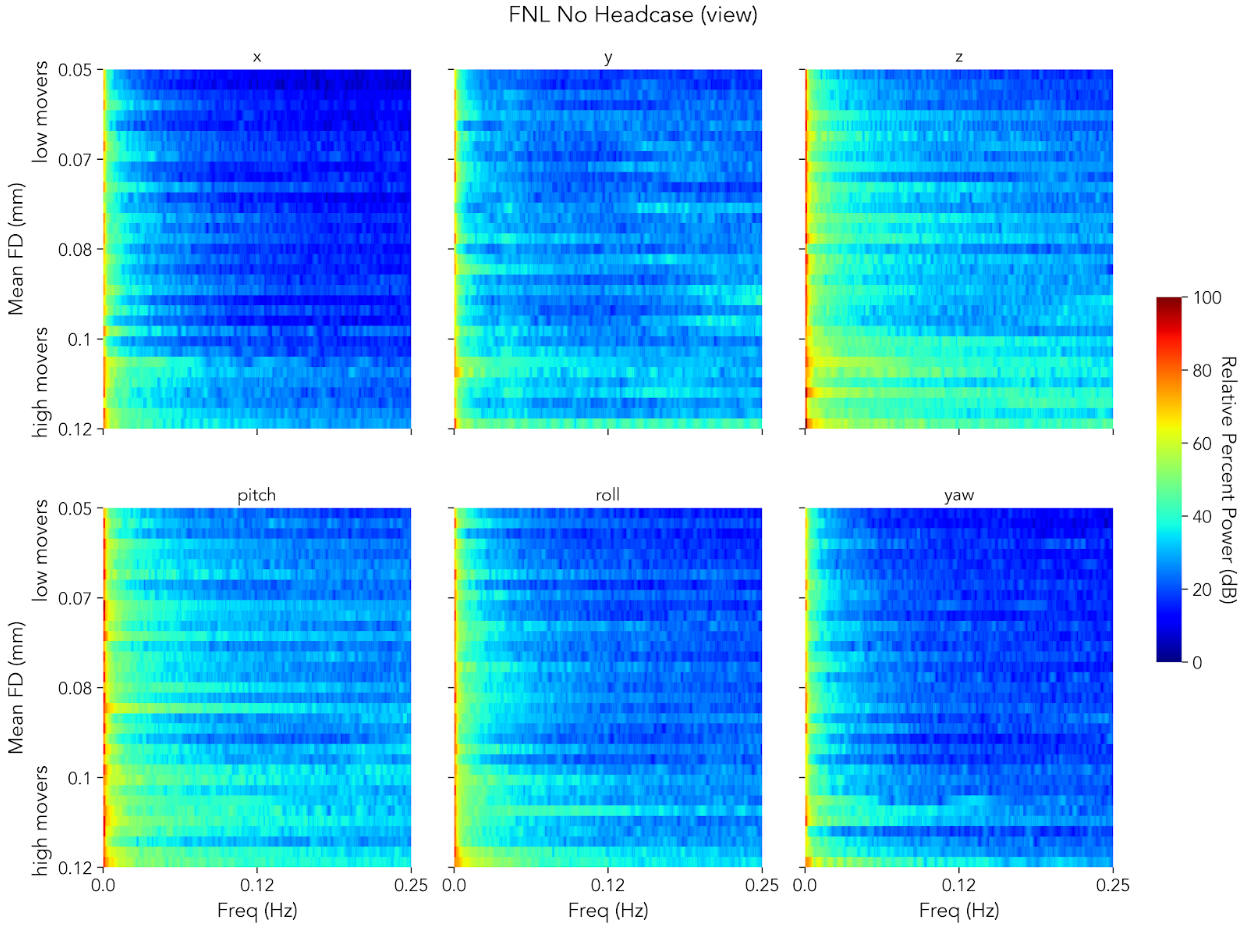

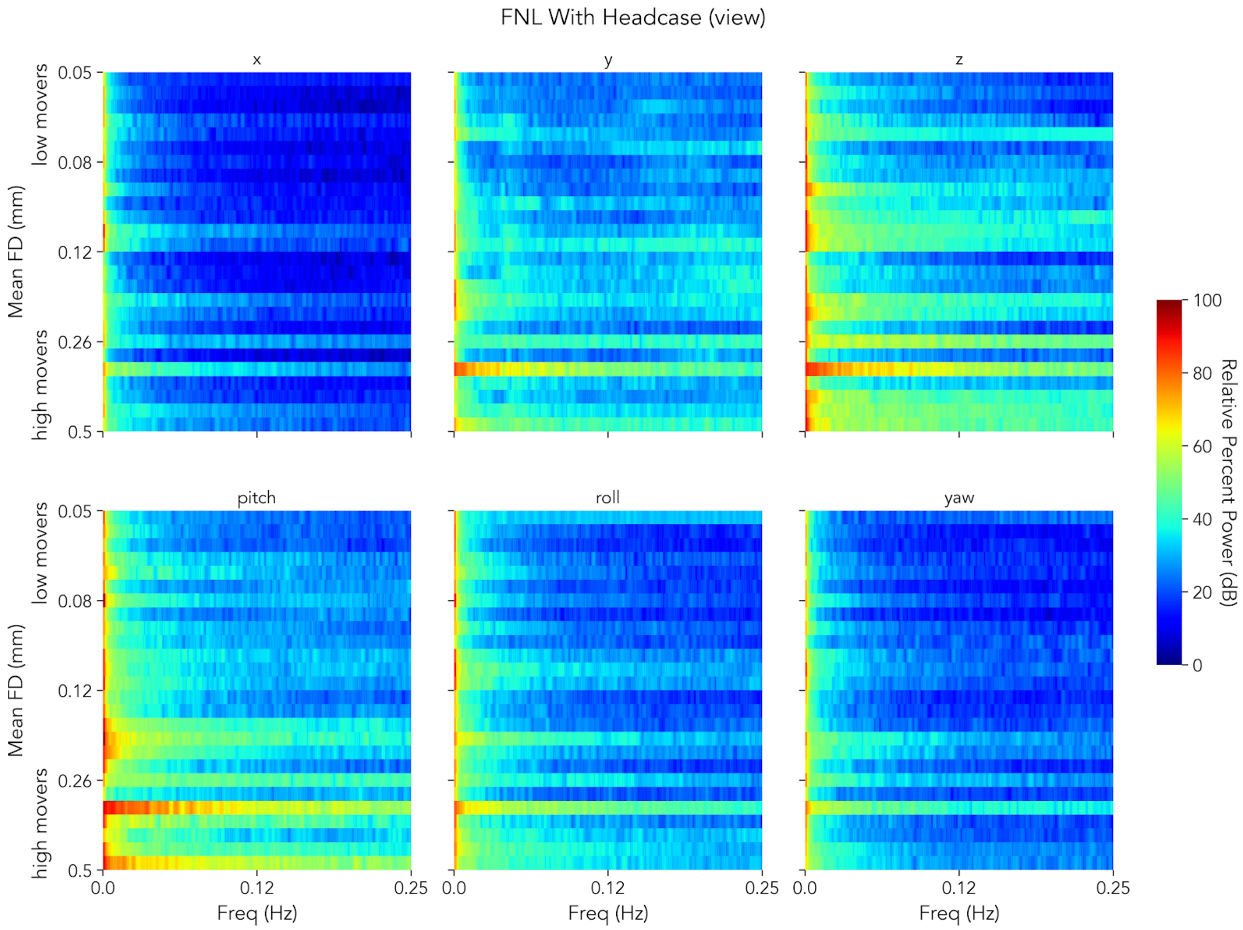

FNL With Headcase (view)
x
y
z
pitch
roll
yaw
low movers
high movers
Mean FD (mm)
0.05
0.08
0.12
0.26
0.5
Freq (Hz)
0.0
0.12
0.25
Relative Percent Power (dB)
100
80
60
40
20
0

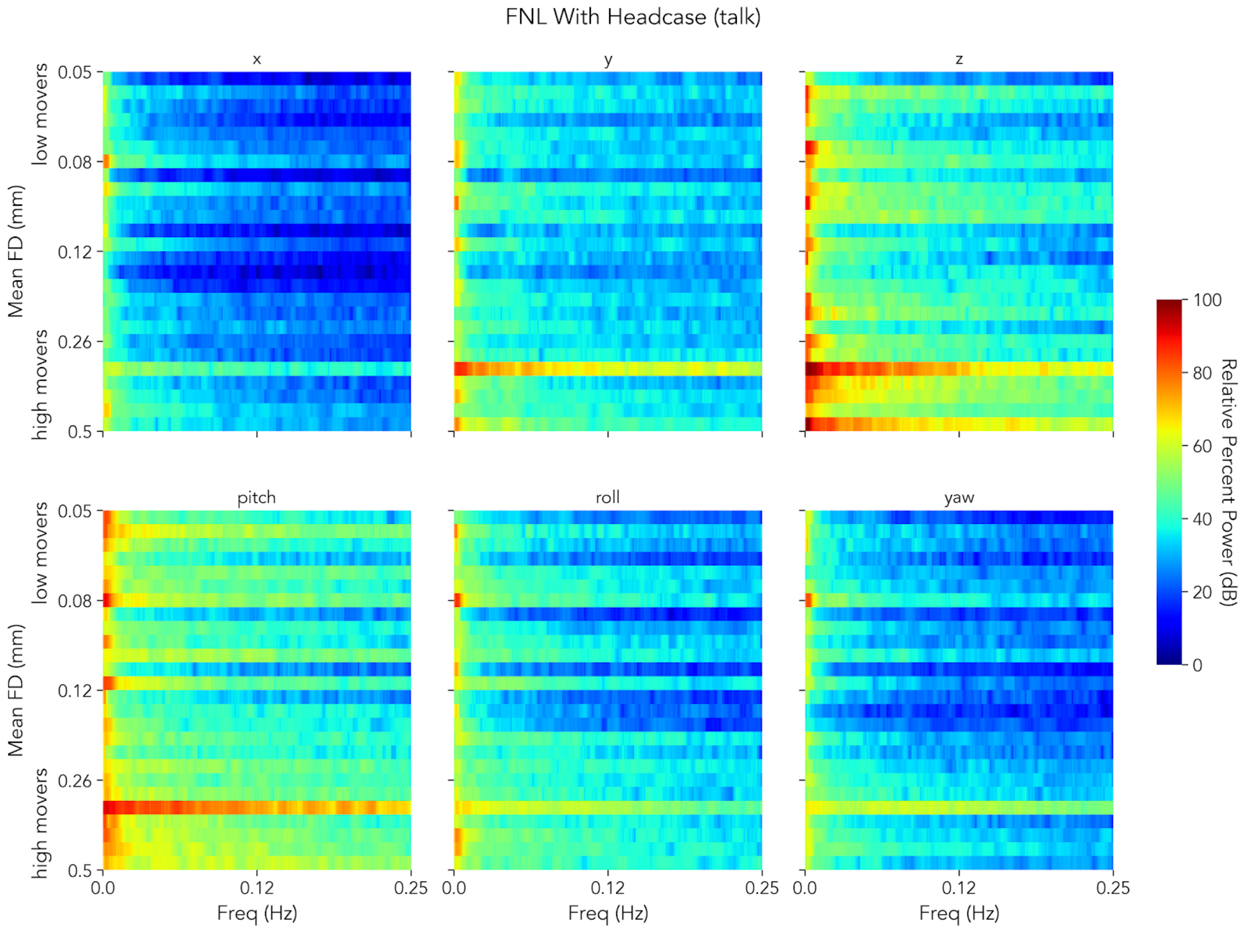

FNL With Headcase (talk)
x
y
z
pitch
roll
yaw
Mean FD (mm)
low movers
high movers
0.05
0.08
0.12
0.26
0.5
0.0
0.12
0.25
Freq (Hz)
Relative Percent Power (dB)
100
80
60
40
20
0

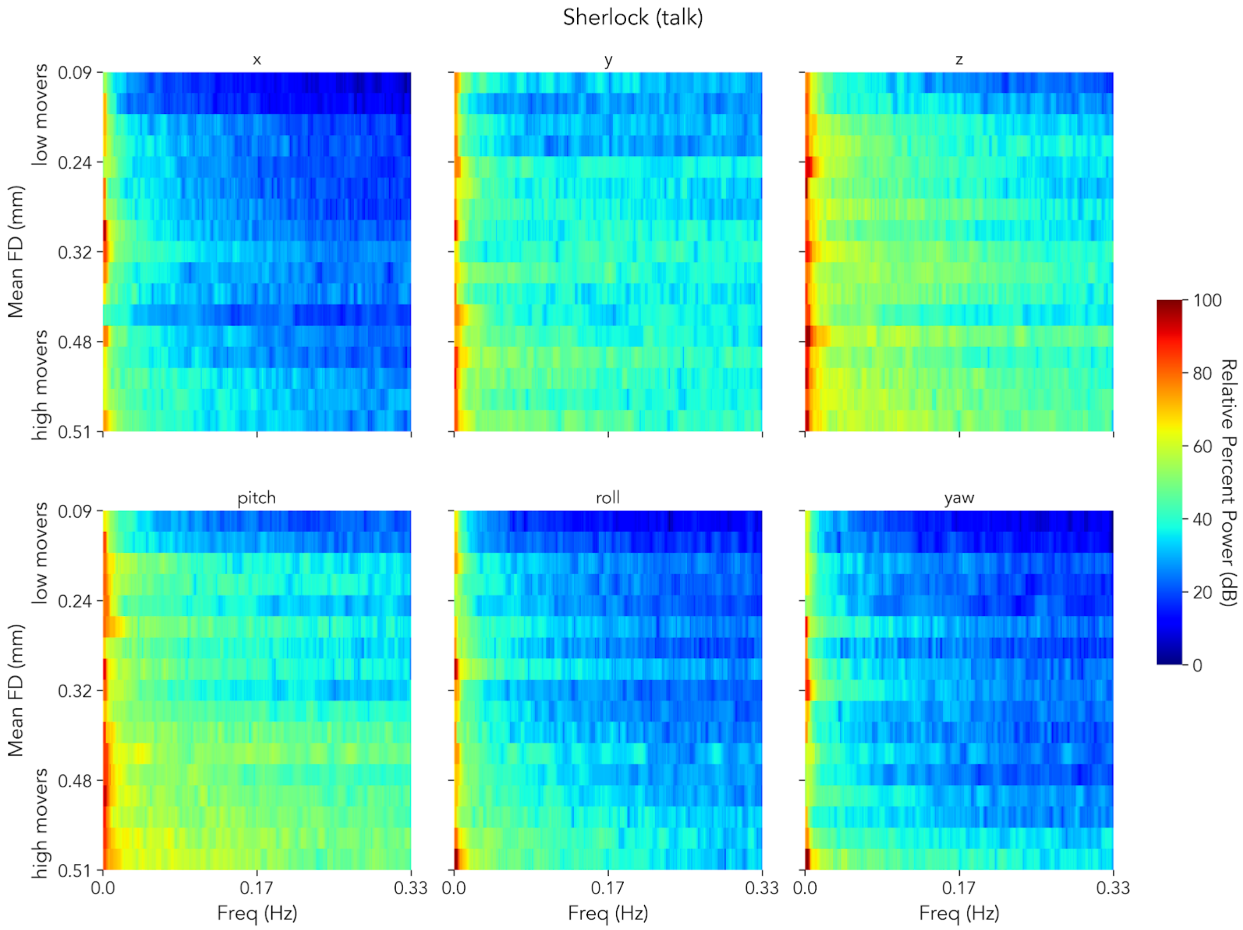
Sherlock (talk)
x
y
z
pitch
roll
yaw
low movers
high movers
Mean FD (mm)
0.09
0.24
0.32
0.48
0.51
0.0
0.17
0.33
Freq (Hz)
Relative Percent Power (dB)
100
80
60
40
20
0

**Table 1. Head case comparisons while *viewing* movies, excluding 2 high motion participants in the FNL with headcase dataset.**

| *Comparison* | *Condition* | *Metric* | *Mean.Difference* | *t* | *p.perm* |
|---|---|---|---|---|---|
| FNL no-case - FNL with-case | Viewing | FD Mean | 0.008 (-0.015 0.031) | 0.689 | 0.496 |
| FNL no-case - FNL with-case | Viewing | FD Median | 0.01 (-0.005 0.024) | 1.309 | 0.196 |
| FNL no-case - FNL with-case | Viewing | Spike Proportion | 0.001 (-0.023 0.025) | 0.115 | 0.906 |
| FNL no-case - Sherlock | Viewing | FD Mean | -0.008 (-0.032 0.015) | -0.703 | 0.489 |
| FNL no-case - Sherlock | Viewing | FD Median | -0.017 (-0.034 -0.002) | -2.061 | **0.045** |
| FNL no-case - Sherlock | Viewing | Spike Proportion | 0.009 (-0.016 0.033) | 0.723 | 0.466 |
| Sherlock - FNL with-case | Viewing | FD Mean | 0.017 (-0.01 0.043) | 1.248 | 0.224 |
| Sherlock - FNL with-case | Viewing | FD Median | 0.027 (0.01 0.045) | 2.981 | **0.005** |
| Sherlock - FNL with-case | Viewing | Spike Proportion | -0.008 (-0.036 0.019) | -0.553 | 0.576 |

**Table 2. Head case comparisons while *viewing* movies after excluding high motion TRs and excluding 2 high motion participants in the FNL with headcase dataset**

| *Comparison* | *Condition* | *Metric* | *Mean.Difference* | *t* | *p.perm* |
|---|---|---|---|---|---|
| FNL no-case - FNL with-case | Viewing | FD $Mean_{Excluded}$ | 0.009 (-0.004 0.022) | 1.391 | 0.167 |
| FNL no-case - FNL with-case | Viewing | FD $Median_{Excluded}$ | 0.01 (-0.003 0.022) | 1.451 | 0.151 |
| FNL no-case - Sherlock | Viewing | FD $Mean_{Excluded}$ | -0.011 (-0.025 0.002) | -1.541 | 0.134 |
| FNL no-case - Sherlock | Viewing | FD $Median_{Excluded}$ | -0.018 (-0.032 -0.004) | -2.368 | 0.022 |
| Sherlock - FNL with-case | Viewing | FD $Mean_{Excluded}$ | 0.02 (0.005 0.036) | 2.581 | **0.014** |
| Sherlock - FNL with-case | Viewing | FD $Median_{Excluded}$ | 0.027 (0.012 0.044) | 3.373 | **0.002** |

**Table 3. Head case comparisons while *talking*, excluding 2 high motion participants in the FNL with headcase dataset. Excluded values reflect comparisons after excluding high motion TRs.**

| *Comparison* | *Condition* | *Metric* | *Mean.Difference* | *t* | *p.perm* |
|---|---|---|---|---|---|
| Sherlock - FNL with-case | Talking | FD Mean | -0.091 (-0.168 -0.014) | -2.250 | **0.037** |
| Sherlock - FNL with-case | Talking | FD Median | -0.068 (-0.13 -0.005) | -2.058 | 0.055 |
| Sherlock - FNL with-case | Talking | Spike Proportion | -0.145 (-0.281 -0.005) | -1.998 | 0.062 |
| Sherlock - FNL with-case | Talking | FD $Mean_{Excluded}$ | 0.002 (-0.016 0.019) | 0.186 | 0.857 |
| Sherlock - FNL with-case | Talking | FD $Median_{Excluded}$ | -0.003 (-0.024 0.017) | -0.272 | 0.791 |
